# Glial ceramide orchestrates Lipid Droplet homeostasis and age-dependent motor function in *Drosophila*

**DOI:** 10.64898/2026.03.01.708786

**Authors:** Lovleen Garg, H Karthik, D Senthilkumar, Anuradha Ratnaparkhi, Girish Ratnaparkhi

**Author notes:** Author for Correspondence, Address: Biology, Indian Institute of Science Education & Research (IISER), Dr. Homi Bhabha Road, Pashan, Pune 411008, India.

## Abstract

Glia cells have emerged as equal partners to neurons in the development and maintenance of the nervous system.

In this study, we examine the roles of ceramide metabolism and intracellular transfer in glia, contrasting these with those of neurons. We find that glia are more sensitive to the cessation of ceramide synthesis than neurons. Knockdown of ER-localised ceramide synthesis enzymes in glia, but not in neurons, leads to age-dependent motor defects. Intriguingly, Lipid Droplet (LD) size and density are lowered when either glial ceramide is reduced or its transfer to the Golgi, by Ceramide Transfer protein (CERT), is disturbed. Glial CERT knockdown or disruption of its interaction with the ER tethering protein VAPB also affects both motor function and LD dynamics, highlighting the importance of targeted, efficient non-vesicular ceramide transfer at membrane contact sites.

Our research implicates reduced flux through the sphingolipid pathway in glial cells as a critical determinant of adult motor function, with LDs serving as a sensitive diagnostic readout. As such, our study has implications for a host of human motor neuron diseases that show late-onset motor deficits.

**Summary Statement:** We find that the production and transfer of ceramides in glial cells are required for both Lipid Droplet formation and maintenance of adult motor function.

## Introduction

Neurons and glia partition essential metabolic roles that are required for the development and maintenance of the nervous system. Glia roles include providing structural stability, supplying nutrients, recycling neurotransmitters and maintaining a secluded, insulated environment for neurons. Glial cells also shoulder the bulk of lipid production and supply, allowing neurons to focus on their role in transmitting electrical and chemical signals (Barber and Raben, 2019; Lee et al., 2021; Wang et al., 2026). Glial cells produce fatty acids, sphingolipids, sterols, and phospholipids. Of these, sphingolipids are bioactive lipids that contain a long-chain sphingoid base linked to a fatty acyl chain via an amide bond. Sphingolipids are quite diverse in terms of different chain lengths, hydroxylation, and saturated bonds of both the base as well as fatty acid moieties (Bikman and Summers, 2011; Islam et al., 2025; Pan et al., 2023). Ceramides serve as the core building block for the formation of the more complex sphingolipids and glycosphingolipids, with ceramide/sphingolipid metabolism essential to maintain cellular homeostasis (Bikman and Summers, 2011; Pant et al., 2020) (Mencarelli and Martinez-Martinez, 2013). Ceramides are not only crucial structural components of the membranes but also act as signaling molecules, regulating diverse physiological processes including apoptosis, autophagy, inflammation and mitochondrial activity (McCluskey et al., 2022; Oswald et al., 2015; Pan et al., 2023; Vos et al., 2021; Wanikawa et al., 2020; Zhu et al., 2025). Ceramide dysregulation is linked to several disorders like type-II diabetes, fatty liver disease and neurodegenerative disorders (Cutler et al., 2002; Hebbar et al., 2015; Henriques et al., 2017; Islam et al., 2025; Jeon et al., 2023; Pant et al., 2020; Pant et al., 2019). Ceramides have important roles in the context of the nervous system, with age-dependent lipid profiling of the brain highlighting their role in both neurodevelopment and maintaining functionality of the adult brain (McCluskey et al., 2022; McInnis et al., 2024; Mencarelli and Martinez-Martinez, 2013; Pan et al., 2023; Theisen et al., 2025; Uranbileg et al., 2024).

Within cells, Ceramides are at the centre of a sphingolipid metabolic network connecting multiple pathways of synthesis, recycling and breakdown. Ceramides are synthesized through a *de novo* pathway that occurs at the cytosolic leaflet of Endoplasmic Reticulum (ER) membrane through a series of enzymes that are conserved across species (Jeon et al., 2023; McCluskey et al., 2022; Mencarelli and Martinez-Martinez, 2013; Pant et al., 2020). A salvage pathway recycles sphingolipids to ceramides at various locations of the cell (Chen et al., 2022; Hebbar et al., 2015; Islam et al., 2025; Pan et al., 2023; Theisen et al., 2025;Vacaru et al., 2009). *Drosophila melanogaster* is also able to generate ceramide from scratch using a similar set of enzymes and substrates (Kraut, 2011; Pan et al., 2023) and unlike vertebrates, usually has one paralog at each step of synthesis. For example, *Drosophila* has a single Ceramide synthase (CERS), *schlank*, as compared to six *CERS* genes in *H. sapiens* (Bauer et al., 2009; Pathak et al., 2018; Voelzmann and Bauer, 2010).

Once formed at the ER membrane, Ceramides are transported to the Golgi through vesicular or non-vesicular means for the production of sphingolipids (Kentaro Hanada, 2003). Vesicular transport generally requires ATP whereas non-vesicular transport occurs with the help of lipid transfer proteins (LTPs) (Kors et al., 2022; Kovacs et al., 2024; Peretti et al., 2008; Weber-Boyvat et al., 2015). One such LTP is Ceramide Transfer protein (CERT1/CERT), a cytosolic protein present at the ER-Golgi interface. CERT transfers ceramide from ER to *trans-*Golgi where sphingomyelin (Ceramide Phosphoethanolamine, CPE in flies) formation takes place. CERT contains a Pleckstrin Homology (PH) domain at its N-terminus, which binds to PI4P present on the Golgi and a START domain at the C terminus that binds to ceramide at ER (Crivelli et al., 2022; Hanada, 2014; Kawano et al., 2006; Peretti et al., 2008; Rao et al., 2007; Yamaji et al., 2008). CERT is also known to be an interactor of VAMP Associated Protein B (VAPB), an ER resident membrane protein. CERT binds to VAPB, and this interaction assists in the transfer of ceramide from the ER (Kawano et al., 2006; Kumagai and Hanada, 2019; Mishra et al., 2024).

Perturbation in ceramide pathway genes can lead to a host of neurological disorders, including Hereditary Sensory and Autonomous Neuropathy 1 (HSANI), Hereditary Spastic Paraplagia (HSP), Amyotrophic Lateral Sclerosis (ALS), and Parkinson’s (Agrawal et al., 2022; Cutler et al., 2002; Hebbar et al., 2015; Hussain et al., 2019; Islam et al., 2025; Liping Wang, 2022; Pan et al., 2023; Pant et al., 2020; Tracey et al., 2021; Vos et al., 2021). Studies also underscore the importance of ceramide production in the development of the brain (Ghosh et al., 2013; Goyal et al., 2019; Oswald et al., 2015). Moreover, glial CPE, also plays a crucial role in the wrapping of neuronal axons and can affect the circadian rhythm of the flies (Chen et al., 2022; Ghosh et al., 2013; Theisen et al., 2025). Alterations in ceramide levels affect ER structure as well, which can thus lead to disruption in ER function, including lipid droplet (LD) formation (Pant et al., 2019; Rao et al., 2014; Robles-Martinez et al., 2025), which originates at the ER membrane. LDs are intracellular storage domains where neutral lipids, such as triglycerides or sterol esters, are maintained within a layer of phospholipids. The phospholipid layer incorporates a set of specialized metabolic and structural proteins, allowing the LD to function as a metabolic hub (Bohnert and Schrul, 2024; Walther and Farese, 2012). LDs are generally formed at the outer leaflet of the ER membrane whenever there is neutral lipid accumulation in the bilayer. Generally, glia are the major hub for LD formation in the brain and LD dynamics are known to be modulated in neurodegenerative disorders (Goodman et al., 2024; Liu et al., 2015; Pennetta and Welte, 2018; Zhang et al., 2025). Interestingly, ceramide-synthesizing enzymes like *schlank* and Infertile crescent (*ifc*), in flies, have been shown to affect LD homeostasis in the larvae (Bauer et al., 2009; Sociale et al., 2018; Zhu et al., 2025). But how this correlates with motor function remains poorly explored.

Jairaj Acharya’s laboratory (Rao et al., 2007) report that CERT null flies have age-dependent motor dysfunction. The absence of *CERT* impairs mitochondrial function, shortening the fly lifespan to 30 days. Lack of ceramide transfer also leads to changes in ER network, ER stress and increased autophagy (Rao et al., 2014). *Drosophila CERT* null embryos eclose normally, whereas mouse embryos die from cardiac failure (Rao et al., 2007; Wang et al., 2009). Recently, *CERT* gain-of-function mutants have been found in human patients as well (Gehin et al., 2023). However, tissue-specific functions of ceramide and CERT have not been explored very well in both mammals and flies. Recent studies suggest a diverse set of functions for ceramide synthesis enzymes in the production of ceramides in neurons vs. glia, but how these changes can influence behavioural outcomes is poorly understood (Theisen et al., 2025).

Our interest in CERT stems from its interaction with Vesicle-Associated Membrane Protein-associated protein B (VAPB). VAPB is an important ER protein that acts to tether the ER membrane with membranes of other cellular organelles, such as mitochondria and Golgi, to form membrane:membrane contact sites (MCS)(James and Kehlenbach, 2021; Kors et al., 2022; Murphy and Levine, 2016; Obara et al., 2024). MCS serve as a physiological niche that regulates cellular function. Modulation or disruption of this niche affects cellular function (Blair et al., 2025; Borgese et al., 2021b; Neefjes and Cabukusta, 2021). In one example, A P56S mutation in VAPB causes ALS (Borgese et al., 2021a; Kamemura et al., 2021; Nishimura et al., 2004). We have modelled ALS8/VAPB in flies, focusing on the *VAPB^P58S^*allele (*VAPB^P56S^* in humans) (Chaplot et al., 2019; Deivasigamani et al., 2014; Ratnaparkhi et al., 2008; Tendulkar et al., 2022; Thulasidharan et al., 2024). *VAPB^P58S^* animals show age-dependent motor dysfunction (Moustaqim-Barrette et al., 2014; Thulasidharan et al., 2024), cytoplasmic VAPB-positive inclusions (Chaplot et al., 2019; Moustaqim-Barrette et al., 2014; Thulasidharan et al., 2024) and brain inflammation (Tendulkar et al., 2022). In the course of our studies, we found a differential role of ceramides in the brain for motor function, which led us to a strong correlation between glial ceramide production and its outward transfer from the ER during LD formation and adult, age-dependent motor function.

## Results

### A reverse genetic screen for motor function in the Ceramide pathway

To establish a relationship between ceramide formation and motor function, we procured RNA interference (RNAi) knockdown (KD) lines targeting enzymes involved in *de novo* ceramide synthesis, based on the genes listed by (Kraut, 2011). The genes targeted, and their functions, are listed in Table 1, while their Bloomington *Drosophila* Stock Centre (BDSC) identifiers are listed in Materials and Methods.

**Table 1:** Genes of the Ceramide pathway used for screening. *Drosophila* genes knocked down to uncover function in glia vs neuron. Human orthologs of fly genes are listed in Fig. 1A.

| Gene (Flybase Computed Gene #) | Gene name/Enzyme name | Function | References |
| --- | --- | --- | --- |
| *spt-I/CG4016* | Serine palmitoyltransferase subunit-I, serine C-palmitoyltransferase | Sphingolipid biosynthetic process | (Ghosh et al., 2013) (Oswald et al., 2015) (Goyal et al., 2019) |
| *lace/CG4162* | Lace, serine C-palmitoyltransferase | Sphingolipid biosynthetic process | (Ghosh et al., 2013) (Goyal et al., 2019) (Theisen et al., 2025) |
| *Kdsr/CG10425* | 3-ketodihydrosphingosine reductase, 3-dehydrosphinganine reductase | Sphingolipid biosynthetic process | (Pan et al., 2023) |
| *schlank/CG3576* | Schlank, Sphingosine N-acyltransferase | Ceramide synthesis and TAG metabolism | (Bauer et al., 2009) |
| *ifc/CG9078* | Infertile crescent, sphingolipid 4-desaturase | Sphingolipid biosynthetic process, negative regulation of lipophagy | (Zhu et al., 2025) |

The schematic in Fig. 1A shows the *de novo* ceramide synthesis pathway at the ER membrane, which is conserved from yeast to mammals as reported in (Pan et al., 2023). We knocked down each gene (Table 1) using a pan-glial driver (*repo-Gal4*), and F1 males were evaluated for motor function using the ‘Startle Induced Negative Geotaxis’ assay, which is described in the Materials & Methods section. The statistical significance of the individual genes from the screening is shown in the summary table (Fig. 1B’) whereas the bar graph of the climbing assay is plotted in Fig. 1B. KD of each gene is expected to reduce the overall flux of ceramides produced by the ER, with the extent of the reduction determined by the KD efficiency and the gene’s individual role. The data (Fig. 1B, B’) suggest that the absence of ceramides in the glia leads to an age-dependent decline in motor function, with *schlank* emerging as the prominent player in the glia where its KD displayed the most substantial loss of climbing ability, followed by *ifc, lace, Kdsr and spt-I*.

**Figure 1:**
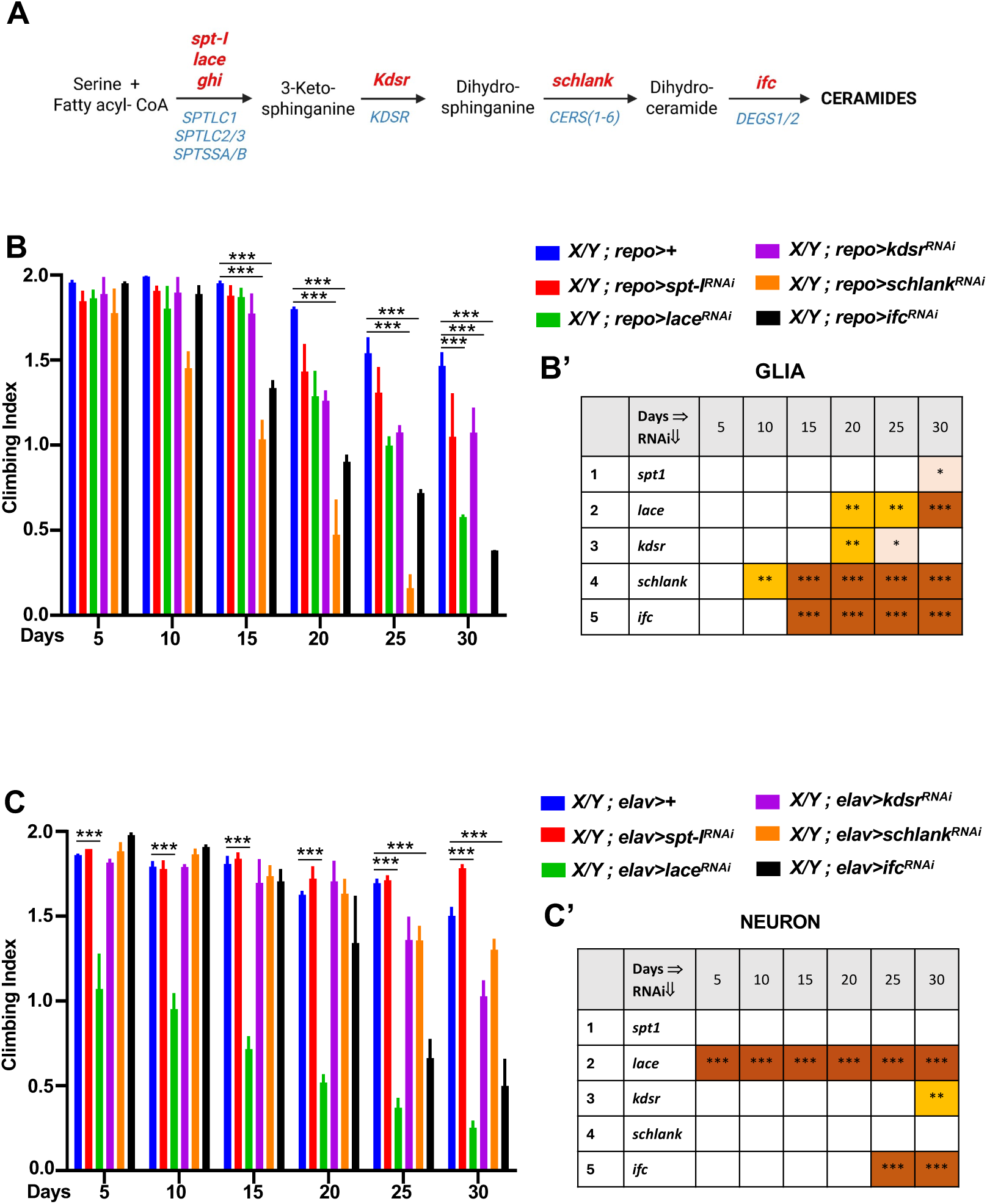
Motor assay screen for enzymes involved in *de novo* ceramide synthesis. (A) Schematic diagram of *de-novo* ceramide synthesis pathway at the ER membrane. *H. sapiens* genes are marked in blue and *D. melanogaster* genes in red. (B) Climbing Index after KD of Glial ceramide synthesis genes in *Drosophila* males, plotted in the form of bar graphs. X-axis represents days post-eclosion (B’) Summary table for climbing assay for male flies with statistical significance shown at each day interval (5, 10, 15, 20, 25, 30) compared to the control. (C) Climbing Index after KD of Neuronal ceramide synthesis genes in *Drosophila* males, plotted in the form of bar graphs. (C’) Summary table for the climbing assay for male flies with statistical significance shown at each day interval (5, 10, 15, 20, 25, 30). n=25-30 males, N=3 biological replicates per genotype. P<0.001(***), <0.001(**) and <0.05(*). Blank spaces in the tables indicate statistical non-significance (ns).

As a counterpoint, we also tested KD of the same set of genes in neurons, using *elav-Gal4* as the driver. *lace* KD in neurons affected motor function the most, as the climbing ability was significantly reduced on day 5 itself, with *ifc* affecting climbing day 25 onwards (Fig. 1C, C’). Intriguingly, *schlank* KD did not affect climbing ability (Fig. 1C). This suggests that the regulation of ceramide dynamics in glia differs from that in neurons. Although these genes have been shown to play differential roles in neurodevelopment (Table 1), this is the first study in the sphingolipid field to report tissue-specific differential roles of ceramides in age-dependent motor function.

### LD homeostasis in Glia is sensitive to ceramide levels

We measured the status of LDs in 5-day adult brains after glial KD of the set of genes (Table 1) involved in ceramide synthesis. LD-specific dye BODIPY493/503 was used to probe LD density and measure LD size/volume, as described in Materials and Methods. We find (Fig. 2A-F) that average LD density decreases on KD of ceramide pathway genes, with *schlank* KD showing the most substantial decrease, followed by *ifc* and *spt-I*. Additionally, *lace* and *Kdsr* KD also showed a slight decline in LD density as well. The data are displayed as bar graphs in Fig. 2G & H. Fig. 2G quantifies LD density (number of LDs per µm^3^) while Fig. 2H shows variation in LD size (Individual LD volume in µm^3^) with KD for all genes leading to a reduction in average LD size and the disappearance of larger LDs, except for the case of *ifc* KD.

**Figure 2:**
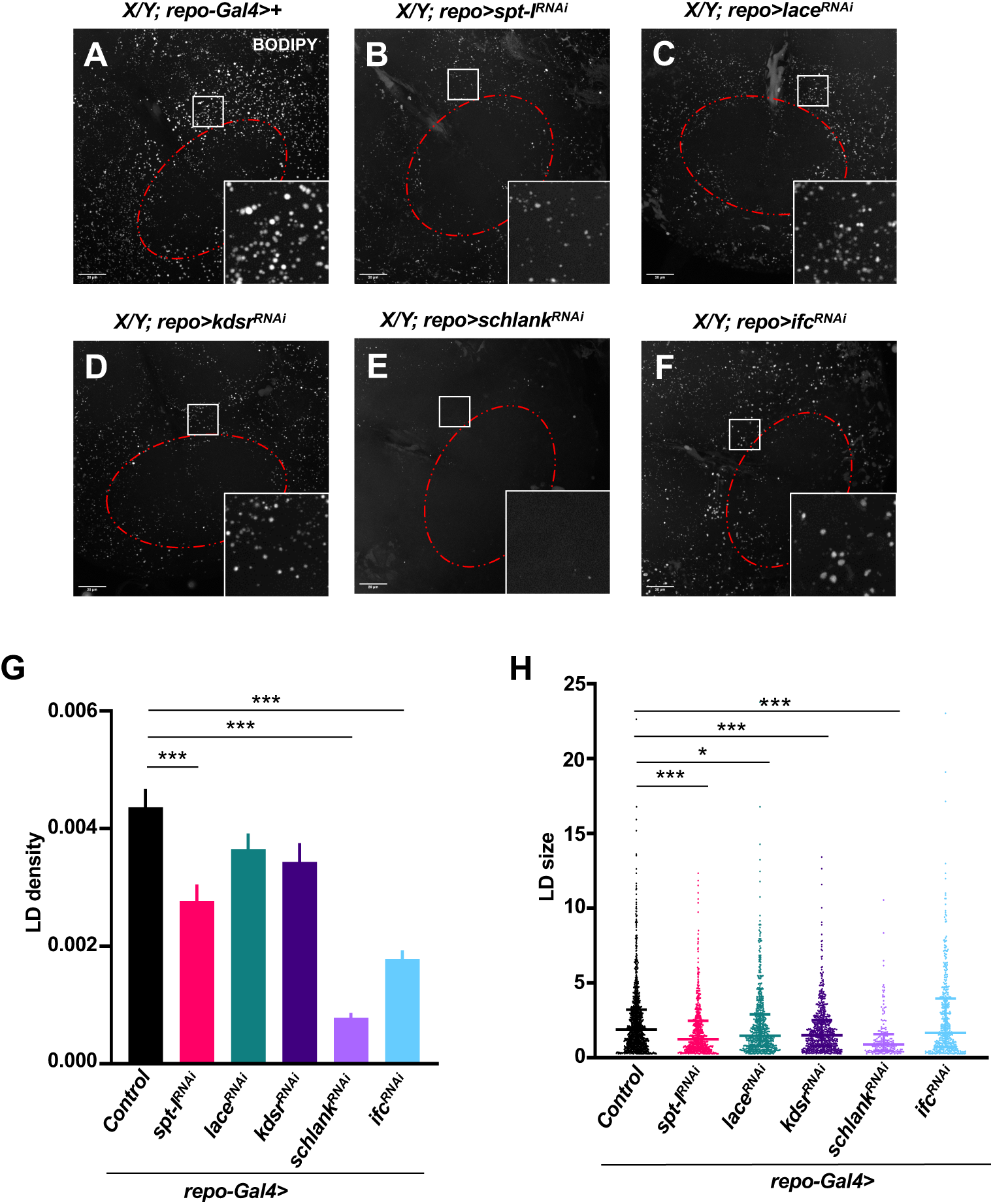
Effect of KD of ceramide pathway genes on glial LD, in the *Drosophila* adult brain. (A-F) represents MIP projections (20% brightness, zoom 1X) of BODIPY-stained LDs in the SEZ (Subesophageal zone) of 5-day-old male adult brains. Glial KD of *de novo* ceramide synthesis pathway genes (B) *spt-I,* (C) *lace,* (D) *Kdsr,* (E) *schlank,* (F) *ifc* showed differential effect on LDs, with *schlank* KD showing the strongest decrease compared to control (A). (G) and (H) represent quantitative analysis for LD density and LD size using five ROIs at the periphery of the SEZ region, marked as a red outline in (A-F). All genes showed a reduction in LD density with *spt-I, schlank,* and *ifc* KD being statistically significant, whereas, in terms of LD size, all genes showed a significant reduction, except for *ifc.* LD density (number of LDs per µm^3^) are plotted as Mean±SEM in the bar graph with statistical significance. For LD size (µm^3^), scattered dot plot with all the lipid droplets from five ROIs of all brain samples from each genotype is represented with median and interquartile range. *p* value is shown as <0.001(***), <0.05(*). n=5 ROIs per brain, N=5-6 adult brains per genotype.

Thus, a decrease in ceramide synthesis in the ER leads to fewer and smaller LDs, and this broadly correlates with age-dependent loss of motor function, with *schlank* KD having the most substantial effect on both LD formation and age-dependent decrease of motor function.

### Lack of Ceramide transfer leads to age-dependent motor dysfunction

In the previous section, we have established that the role of ceramide synthesis in the glia is essential for motor function as well as LD homeostasis. We were interested in whether ceramide transfer from the ER, by CERT, post-synthesis, also affects these functions. A schematic of CERT function (Fig. 3A) and its domain structure (Fig. 3B) highlights its location in the MCS and its interaction with the ER-based MCS architecture protein VAPB (Kumagai and Hanada, 2019; Mishra et al., 2024; Perry and Ridgway, 2006; Slee and Levine, 2019). The Acharya lab generated a CERT null line, *dCERT^1^* (Rao et al., 2007) and were the first to demonstrate that these mutants had reduced lifespan, metabolic dysfunction, reduced thermal tolerance, and locomotor defects (Rao et al., 2007). (Gehin et al., 2023) also reported loss of motor activity because of a lack of *CERT* function in flies. Here, we first reproduced the *dCERT^1^* (called *CERT^∆^* henceforth) motor phenotype, confirming that *CERT^∆^* homozygotes cannot climb efficiently in ‘startle-induced negative geotaxis assays’, showing a reduced climbing index on day 5, with further decline with age. At day 25, *CERT^∆^* animals could not climb at all. The *CERT* allele was haplosufficient for climbing ability, with *CERT^∆^*/+ animals showing no reduction in climbing ability over 25 days of ageing. *CERT^∆^*/Df animals had a stronger climbing phenotype and shorter lifespan, dying by day 15 (Fig. 3C).

**Figure 3:**
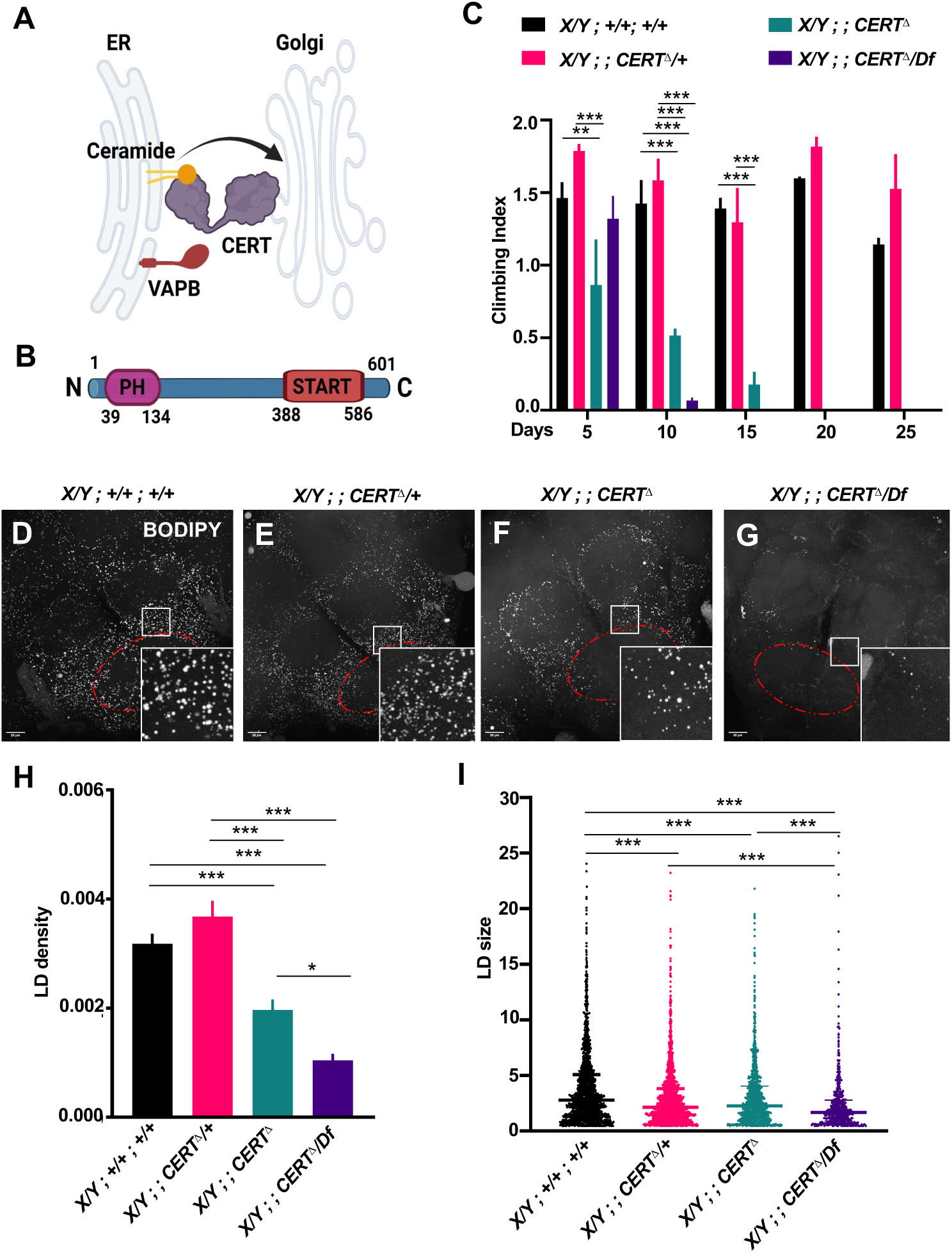
*CERT* loss in *Drosophila* affects motor function and LD homeostasis. (A) Schematic representation of ceramide transfer from ER to *trans-*Golgi mediated by CERT with the help of ER resident VAPB. (B) Domain architecture of *Drosophila* CERT (C) Homozygous *CERT* null mutant (*CERT^Δ^*) exhibit significant motor defects at day 5 itself, which worsen with age. However, the heterozygous mutant climbs at par with wild-type flies. *CERT^Δ^*/Df showed progressive motor impairment and completely stopped climbing after day 10. Mean±SEM values are plotted in the graph representing climbing index on Y-axis with number of days on X-axis with statistical significance as p<0.001(***). n=25-30 males, N=3 replicates per genotype. (D-G) shows representative MIPs (40% brightness) of BODIPY-stained LDs in 5-day-old male adult brains. *CERT^Δ^* (F) and *CERT^Δ^/Df* (G) have fewer LDs compared to wildtype (D) and *dCERT^Δ^*/+ (E). (H) and (I) shows quantitative analysis of LDs with LD density and LD size as parameters analysed using five ROIs around SEZ region (depicted with red outline in panel D). n=5 ROIs, N=5 adult brains. For LD density (number of LDs per µm^3^), Mean±SEM values are plotted in the graph with p<0.001(***), <0.01(**), <0.05(*). For LD size (µm^3^), scattered dot plot with all the lipid droplets from five ROIs is represented with median and interquartile range and p<0.001(***), <0.01(**), <0.05(*).

### Lack of Ceramide transfer perturbs LD homeostasis

The *CERT^∆^* line showed an age-dependent loss of motor function (Fig. 3C), with 30% climbing ability lost on Day 5 post eclosion (Fig. 3C, Green Bar). We were interested in whether ceramide transfer can also affect LD dynamics. On probing LDs for *CERT^∆^* mutant brain samples at day 5, we found wild-type LD density was 0.0032/µm^3^ and for *CERT^∆^*/+, 0.0037/µm^3^, indicating that *CERT* is haplo-sufficient in terms of LD density (Fig. 3D-G). Interestingly, LD density dropped by 40% to 0.002/µm^3^ on day 5 in *CERT^∆^* adult brains (Fig. 3G). *CERT^∆^/Df* showed a 60% decrease in LD density to 0.001/µm^3^ (Fig. 3G-H), validating the reduction in overall LD density in the absence of CERT. Both *CERT^∆^* and *CERT^∆^/Df* showed a reduced average LD size, with *CERT^∆^/Df* having the lowest median LD size in this experimental set. Intriguingly, *CERT^∆^* and *CERT^∆^*/+ had similar average LD size, suggesting that LD size determination requires a full CERT allelic dose (Fig. 3I).

### Lack of glial *CERT* leads to age-dependent motor defects in *Drosophila*

Earlier (Fig. 1) we uncovered a correlation between glial ceramide production and motor function. CERT null animals (Fig. 3) also showed age-dependent loss of climbing ability. To dissect tissue-specific roles for CERT, we both overexpressed (OE) and KD *CERT* via RNAi using a set of tissue-specific drivers and subsequently evaluated climbing ability in the adult F1 fly over 25-30 days post-eclosion. For OE, we generated a transgenic *UAS-CERT* line, as described in the Materials & Methods section. First, we used a ubiquitous driver, *tubulin-Gal4* (*tub-Gal4*), to modulate *CERT* and assessed fly motor activity. The ubiquitous *CERT* KD (Fig. 4A, A’) recapitulated, though to a lesser extent, the whole-body loss-of-function (Fig. 3C), confirming that loss of *CERT* led to age-dependent motor deterioration. *CERT* OE did not lead to climbing defects, with a mild decrease in climbing index seen on day 25.

**Figure 4:**
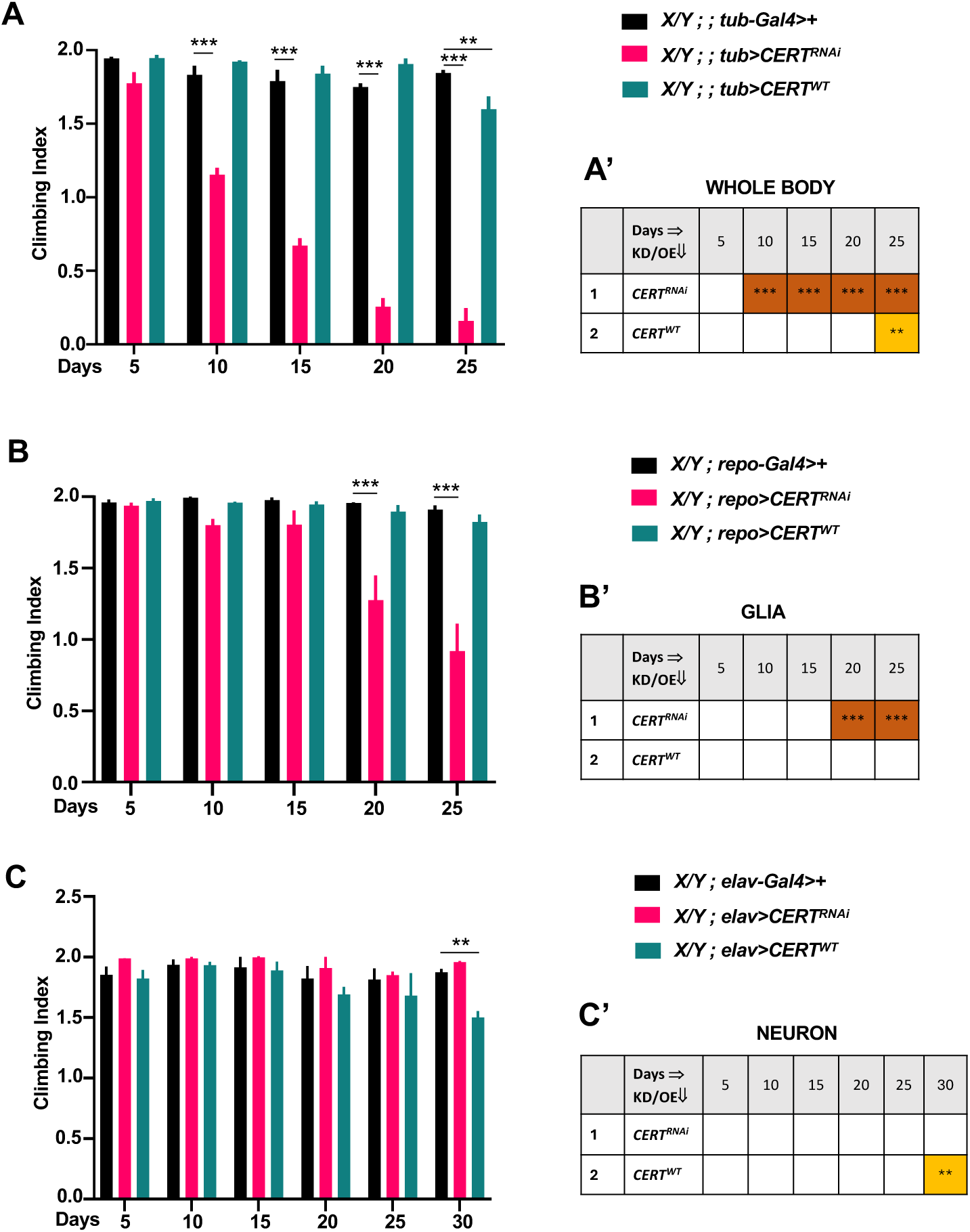
Tissue-specific *CERT* role in motor function. (A) Climbing Index is plotted for whole body modulation of *CERT* using *tub-Gal4* in male flies with statistical significance. *CERT KD* showed progressive motor defects with age, whereas OE caused mild defects observed at day 25. (A’) Summary table for statistical significance for each day interval (5, 10, 15, 20, 25, 30) compared to control. (B) Climbing Index is plotted for glial modulation of *CERT* using *repo-Gal4* in male flies with statistical significance. Glial KD of *CERT* in *Drosophila* led to motor defects by day 20 and kept on worsening with age, whereas *CERT* OE did not cause any detectable motor impairment. (B’) Summary table for statistical significance for each day interval (5, 10, 15, 20, 25, 30) compared to the control. (C) Climbing Index is plotted for neuronal modulation of *CERT* using *elav-Gal4* in male flies with statistical significance. Neuronal KD of *CERT* showed no defect in motor activity whereas overexpressing *CERT* led to mild motor defects on day 25. (C’) Summary table for statistical significance for each day interval (5, 10, 15, 20, 25, 30) compared to control. Mean±SEM values are plotted in the graph. p<0.001(***), <0.01(**). n=25-30 males, N=3 replicates per genotype.

Next, in order to test the tissue-specificity of CERT function, we OE and KD *CERT* in both glia (*repo-Gal4*) and neurons (*elav-Gal4*) (Fig. 4). Interestingly, glial KD led to a loss of climbing ability from day 20 onward, whereas neuronal knockdown did not. OE of *CERT* did not show any climbing defects over age, with a mild reduction in the climbing index for neuronal OE on day 30 (Fig. 4B-C).

Our data suggests that the transfer of glial ceramide by CERT from the ER is important for the animal to maintain its motor function, viz its climbing ability, with the neuronal role being less important, vis-à-vis motor function. We also used an alternative neuronal driver, *nSyb-Gal4,* to KD *CERT* in glia (Suppl. Fig. 1). Here, again, we see that climbing ability is normal till day 20, and *CERT-KD* and OE flies start showing major motor defects after day 25 onwards.

### Lack of glial ceramide transfer leads to LD defects in the adult brain

Since ceramide transfer by glial CERT is important for motor function, does *CERT* KD also affect LD dynamics? *CERT* KD in glia (Fig. 5A-C, G-H) led to a ∼60% reduction in LD density. The drop was stronger when a more efficient Valium 20 RNAi line was used rather than the maternal Valium22 RNAi line (Mishra et al., 2024). The reduction in LD density was of the same order as seen in *schlank* (80%) and *ifc* (50%) KDs (Fig. 2) as well as *CERT^Δ^/Df* (Fig. 3H). *CERT* KD also led to a change in the distribution of LD size, with larger LDs not observed, much like in the case of *schlank* KD and *CERT^Δ^/Df* (Fig. 2H, Fig. 3I).

**Figure 5:**
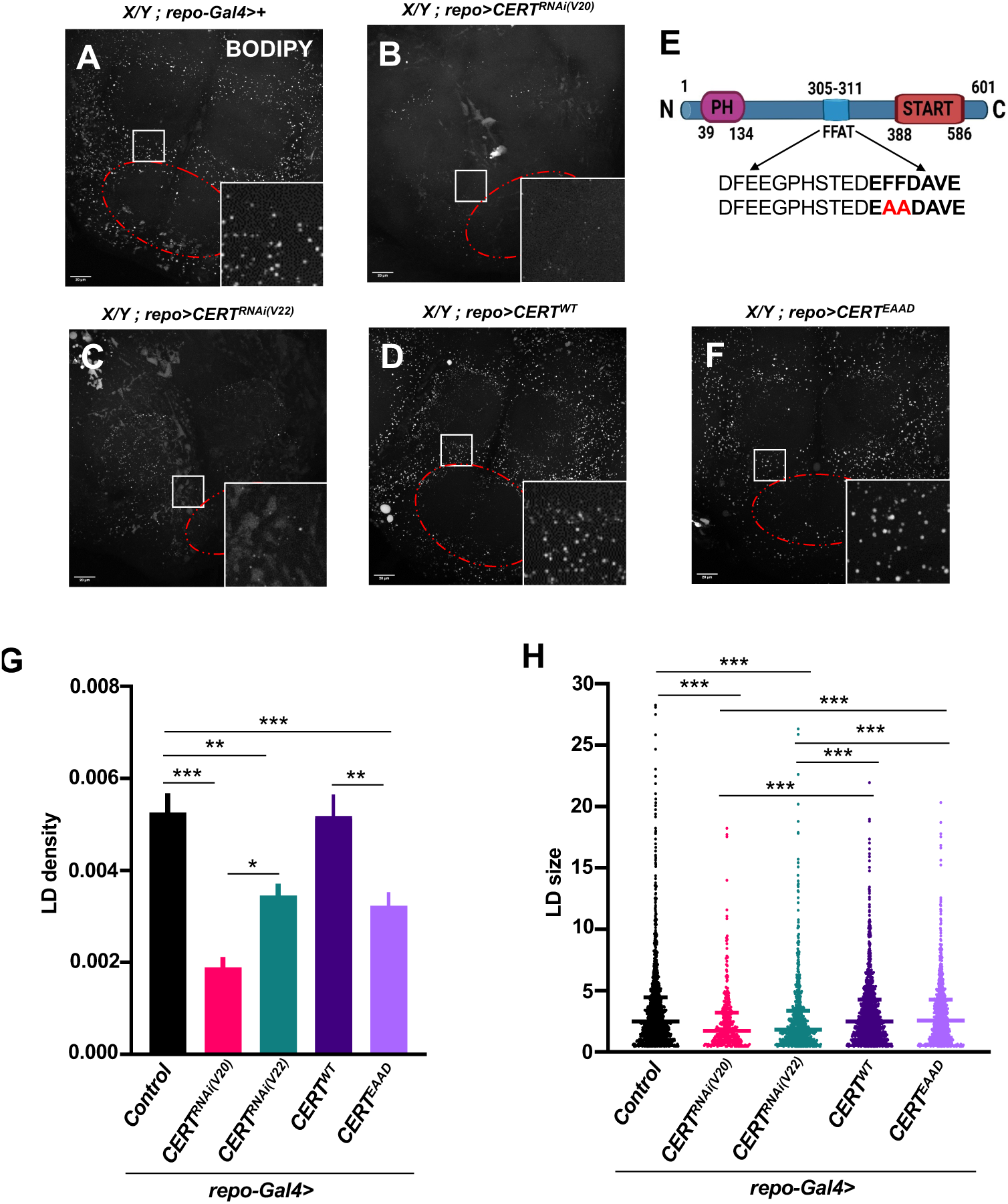
Glial *CERT* affects LD dynamics in the *Drosophila* adult brain. (A-D, F) Representative MIPs (40% brightness) of LDs stained with BODIPY in 5-day-old male adult brains. *CERT* KD in glia using Valium 20 (B) and Valium 22 (C) RNAi lines led to fewer LDs with a smaller size compared to control (A). However, *CERT^WT^* OE (D) showed no change at all compared to the control. (E) Schematic representation of the FFAT motif present in CERT, and the mutation (FF>AA) perturbing the binding motif. (F) *CERT^EAAD^* OE shows a significant reduction in LDs but no change in size. (G) and (H) Quantitative analysis of LDs with LD density and LD size as parameters analyzed using five ROIs around SEZ region (depicted with red outline in panel A). Mean±SEM values are plotted for LD density (number of LDs per µm^3^) in the graph. For LD size (µm^3^), scattered dot plot with all the individual lipid droplets from five ROIs of all brain samples is represented with median and interquartile range. Statistical significance is shown as p<0.001(***), <0.01(**), <0.05(*). n=5 ROIs per adult brain, N=5 brains for each genotype.

Excess *CERT* did not appear to affect LD dynamics, with both LD density and size in the brain similar to wild-type (Fig. 5D, G & H). Since CERT is physical interactor of VAPB in flies as well (Mishra et al., 2024) and VAPB is known to interact with a lot of proteins via their canonical FFAT motif we tested the role of CERT’s interaction with VAPB and any role for the interaction in modulating LD density and size. CERT has conserved FFAT motif (EFFDAVE, which is important for ceramide transfer (Kawano et al., 2006)). We generated a transgenic CERT line (*CERT^EAAD^*) with the FFAT motif modified from (FF to AA) to disrupt its interaction with VAPB (Fig. 5E). Interestingly, male flies overexpressing CERT^EAAD^ in both neurons and glia (Suppl. Fig. 1 & 2) did not show any climbing defects, although flies were sluggish in the case of glia OE. Intriguingly, glial OE of the *CERT^EAAD^* line led to a significant reduction in LD density but no change in LD size (Fig. 5F-H). This suggests that an excess of CERT that does not bind VAPB efficiently can perturb LD density, but not LD size.

### Motor function can be rescued by transgenic glial CERT

(Gehin et al., 2023) generated transgenic *CERT^WT^* and *CERT^SL^* lines expressed under genomic promotors referred as *gCERT^WT^* and *gCERT^SL^*. *gCERT^SL^* corresponds to expression of *CERT* with Ser to Leu mutation at 149^th^ position (132^th^ position in humans). *gCERT^SL^* is a hyperactive allele which shows CerTra syndrome-like phenotypes in flies. Briefly, *CERT* gain of function led to the production of more ceramides, dihydroceramide, CPE, as well as di-CPE in the heads of the flies (Gehin et al., 2023). Interestingly, both genotypes do not show any climbing defects under normal conditions but have climbing defects under mechanical stress. We find that the climbing index is substantially improved when we balance either *gCERT^WT^*and *gCERT^SL^* with *CERT^∆^* (Fig. 6A). The rescue improves the climbing index two-fold when compared to *CERT^∆^,* up to day 20, but does not achieve the climbing index of wild-type flies. Intriguingly, glial OE of *CERT^WT^* (Fig. 6B) shows a more substantial rescue of motor function, with the climbing index being at par with wild-type flies on days 5 and 10, and a 3-fold improvement in climbing index in older flies (days 15, 20 and 25). OE of *CERT^EAAD^* also rescues motor function in an age-dependent manner, with the rescue at par with wild-type CER*T^WT^*(Fig. 6B) for days 5, 10 and 15, with a slight fall in average climbing index on days 20 and 25.

**Figure 6:**
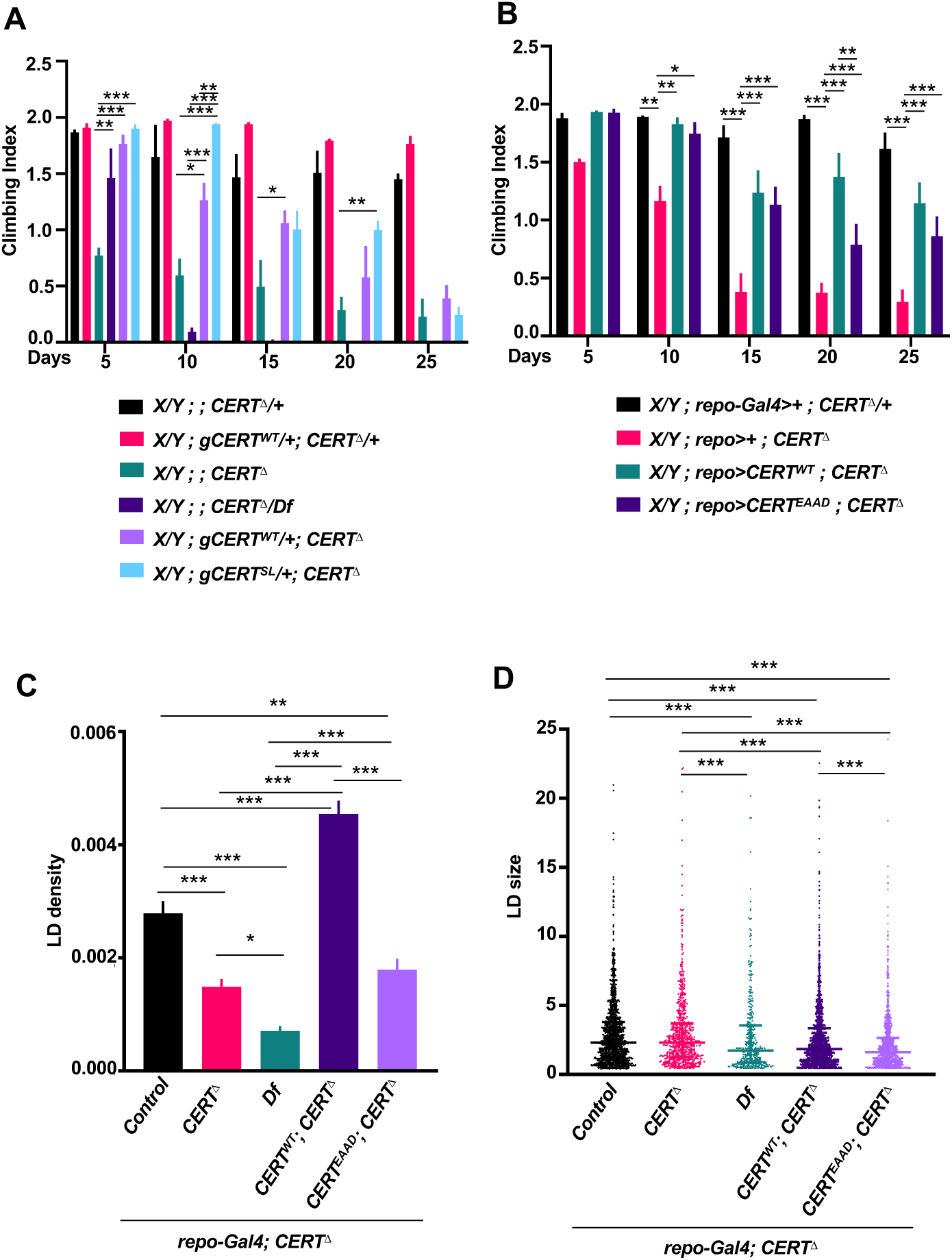
Glial expression of *CERT* rescues motor and LD defects seen in *CERT^Δ^*. (A) Introducing a genomic copy of *CERT^WT^* in *CERT^Δ^* resulted in partial rescue of motor activity, and it behaved completely normally in a heterozygous *CERT^Δ^* mutant. Adding a genomic copy of the hyperactive allele *CERT^SL^*in the *CERT^Δ^* background showed stronger rescue than *CERT^WT^*. (B) Glia-specific expression of *CERT^WT^* in the background of *CERT^Δ^* rescued motor defects significantly, as well as *CERT^EAAD^*. Mean±SEM values of climbing index are plotted in the graph with p<0.001(***) <0.01(**), <0.05(*). n=25-30 males, N=3 biological replicates per genotype. (C-D) Quantitative analysis of LD density (number of LDs per µm^3^) and LD size (µm^3^) were performed on five ROIs surrounding SEZ region. For LD density, data is represented as Mean±SEM in the graph with statistical significance as p<0.001(***). LD size is shown as scattered dot plot with all the lipid droplets from five ROIs of all brain samples of each genotype p<0.001(***). n= 5 ROIs, N=5 adult brains per genotype.

### LD dynamics can be rescued by glial expression of CERT

Both ceramide synthesis and transfer appear to be important in the glia for motor function. Next, we expressed *CERT^WT^* in the glia of 5-day-old *CERT^Δ^* animals (Fig. 6C, Suppl. Fig. 3) to examine if glial expression of *CERT^WT^* could rescue LD dynamics. Intriguingly, LD density was completely rescued in the presence of excess glial CERT (Fig. 6 C-D), however, the *CERT^EAAD^* was not as efficient in rescuing in the *repo* domain, with its overexpression not rescuing either LD density or size significantly (Suppl. Fig. 3E) (Fig. 6 C-D).

## Discussion

Motor function in animals is regulated by the neuromuscular system. The brain plans, initiates and coordinates voluntary movements using specific neuronal circuits with motor neurons carrying signals to the muscles that execute the action. Feedback systems fine-tune motor actions. At the cellular level, motor function is affected when neurons, their glial partners, or muscle cells lose their ability to perform a function. Loss of motor function is seen in motor neuron diseases (MNDs), which are often progressive. These diseases include, but are not limited to ALS, Progressive bulbar palsy, Primary lateral sclerosis and Progressive muscular atrophy (Fullam and Statland, 2021; Gunay et al., 2021; Ilieva et al., 2009; Nowakowski et al., 2025; Rachakonda et al., 2004; Tendulkar et al., 2022; Verma et al., 2025).

Lipid metabolism is central to cellular homeostasis; amongst its diverse roles, it maintains cellular membranes and their fluidity, stores energy, forms signalling networks, and acts as an insulating entity (Bikman and Summers, 2011; Cockcroft and Raghu, 2018; Gunay et al., 2021; Hussain et al., 2019; Islam et al., 2025; Kraut, 2011; Mesmin et al., 2019; Schmitt et al., 2014; Toprak et al., 2020; Walther and Farese, 2012; Yang et al., 2022). Lipid balance, whether in neurons, glia, or muscles, is a major factor in maintaining motor function. Increasingly, MNDs are linked to changes in lipid homeostasis, with lipid dynamics significantly altered in disease; phospholipids, ceramides, and cholesterol show major alterations and serve as biomarkers of disease progression (Agrawal et al., 2022; Byrns et al., 2024; Cutler et al., 2002; Henriques et al., 2017; Hussain et al., 2019; Liu et al., 2020; McCluskey et al., 2022; Oswald et al., 2015; Pan et al., 2023; Pant et al., 2020; Schmitt et al., 2014; Tracey et al., 2021; Vos et al., 2021; Wanikawa et al., 2020; Yang et al., 2022). One major pathway that is altered in MNDs is the Ceramide pathway. Dysregulation of Ceramide metabolism has been linked to neurodegeneration (Pan et al., 2024) (Kolter and Sandhoff, 1999; Pan et al., 2023; Peng et al., 2025). Ceramides are synthesised in the ER and subsequently transferred and distributed across the cell, with the basic ceramide scaffold serving as a substrate for modification to generate a variety of complex ceramide derivatives. An increase or decrease in ceramide species can lead to mitochondrial dysfunction, ER stress, and inflammation, and can result in neuronal or glial cell death. This makes the various enzymes in the ceramide pathway important therapeutic targets, as each enzyme works to convert one lipid to another, maintaining levels across the ceramide/sphingolipid metabolic network (Agrawal et al., 2022; Alessenko and Albi, 2020; Arboleda et al., 2009; Cutler et al., 2002; Goyal et al., 2019; Hebbar et al., 2015; Henriques et al., 2017; Islam et al., 2025; Jung et al., 2017; McCluskey et al., 2022; Pan et al., 2023; Pant et al., 2019; Vos et al., 2021).

Our work highlights several important features of motor neuron function and dysfunction. First, from our screen of ceramide pathway enzymes, it is clear that Neurons and Glia differ significantly in their ability to handle ceramide (Fig. 7). This is supported by (McInnis et al., 2024), and highlighted in recent reviews (Ayub et al., 2021; Islam et al., 2025); who report differential levels of ceramide species in neurons and glia, which may be linked to the animal’s motor function. In our study, Glia appear to be more sensitive to reductions in both ceramide and complex sphingolipids, as alterations at multiple steps in the ceramide pathway lead to motor dysfunction in adult flies.

**Figure 7:**
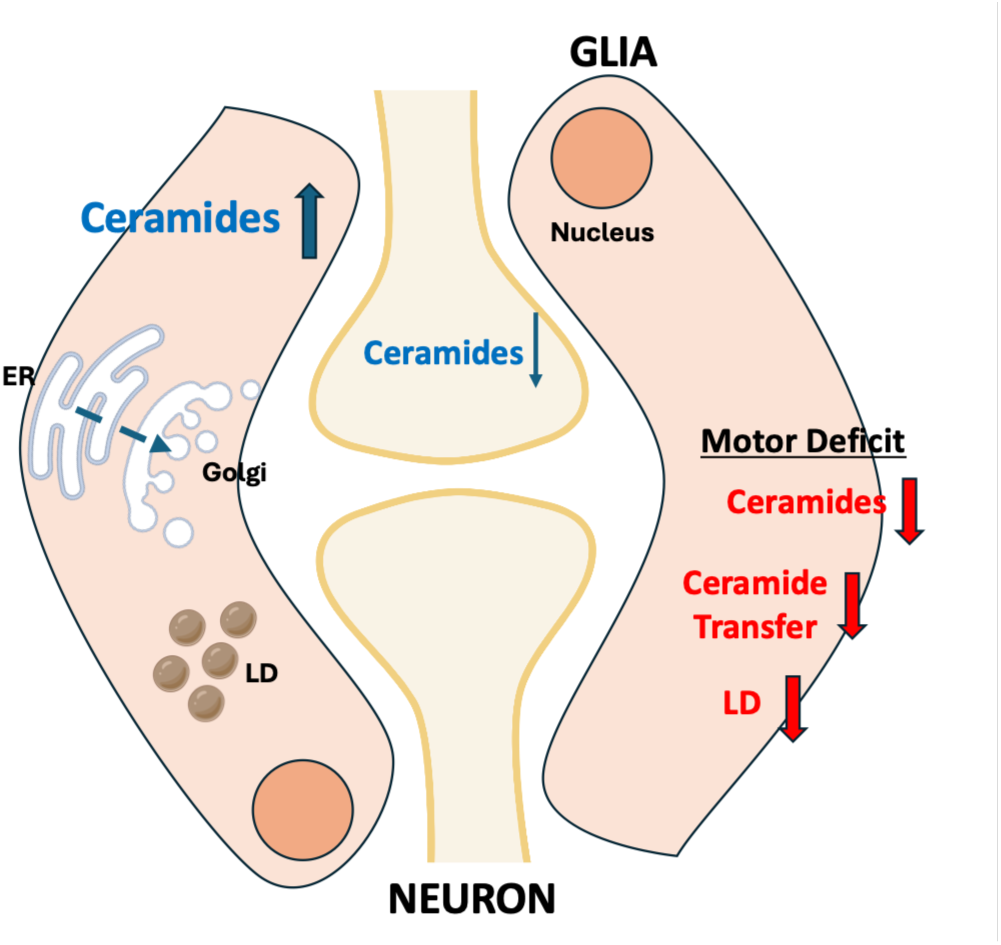
Glial sphingolipid metabolism affects LDs and age-dependent Motor Function. Our current understanding of differential roles for the Ceramide pathway in glia *vs* neurons suggests that the flux of ceramide synthesis from the ER to the Golgi is higher in glial cells, especially those with higher membrane turnover. In contrast, neurons appear to require more complex signaling sphingolipids e.g., Sphingosine-1-phosphate, for function, while keeping basal Ceramide synthesis in check. In glia, we find that a drop in glial Ceramide synthesis or a cessation of its transfer to the Golgi, both of which are required for the generation of complex sphingolipids, leads to motor neuron deficits. The reduction of flux through the ceramide pathway also affects LD dynamics, suggesting cross-communication between the LD and sphingolipid pathways.

Interestingly, in our experiments, *schlank* KD in neurons, which would lead to the accumulation of dihydrosphinganine or reduction in dihydroceramides, did not have any effect on motor function. In contrast, reduction of dihydrosphinganine following neuronal knockdown of *lace* results in severe motor neuron dysfunction. Several studies report differential roles of *spt-I, lace, schlank* and *ifc* in neurodevelopment, with neuronal knockdown of *spt-I* and *lace* affecting mushroom body development, suggesting that sphinganine formation is essential for neuronal function (Goyal et al., 2019; Oswald et al., 2015). Moreover, fatty acids accumulate in neurons in the absence of these enzymes, leading to oxidative stress (Liu et al., 2017; Tang et al., 2024). In the case of *schlank* and *ifc,* studies point to their developmental roles in glia (Chen et al., 2022; Ghosh et al., 2013; Islam et al., 2025; Theisen et al., 2025; Zhu et al., 2025) as the nervous system forms in the animal. We find that in the case of *ifc*, loss in both neurons and glia affects motor function, suggesting that dihydroceramide levels have a toxic impact on both cell types. Interestingly, glial loss of *ifc is* more potent than in neurons, as seen in our motor assay screen and recently reported by (Zhu et al., 2025).

Second, *CERT* KD in glia alone approaches, in terms of penetrance, the age-dependent motor dysfunction seen in whole-body KD or a CERT null animal. This suggests a more important role for CERT in glia than in neurons (Fig. 7), especially in the context of adult motor function. Broadly, our data indicate that the generation of complex sphingolipids (ceramide phosphoethanolamine/sphingomyelin, glycosphingolipids), which are biochemical steps downstream of ER-based ceramide production and transfer, is a critical requirement for motor function. The absence or reduction of these species perturbs the glial-neuronal metabolic equilibrium and disrupts the motor circuit (Fig. 7). This thesis is supported by recent studies (Byrns et al., 2024; Chen et al., 2022; Islam et al., 2025; Liping Wang, 2022; Theisen et al., 2025; Vaughen et al., 2022; Zhu et al., 2025).

Third, perturbing both ceramide production and their transfer affects LD dynamics. This finding is intriguing, as LD formation has no direct relationship with the ceramide synthesis pathway aside from the fact that both processes occur at the ER membrane (Bikman and Summers, 2011; Walther and Farese, 2012). Recently, LDs have been shown to contain acylceramides and small amounts of dihydroceramides (Robles-Martinez et al., 2025; Senkal et al., 2017). Consistent with this, ceramide production or its transfer from the ER results in smaller, less dense LDs. *A priori*, LD formation would depend on the availability of TAG, which in turn would depend on fatty acid availability and the normal functioning of TAG-generating enzymes, namely Glycerol-3-phosphate Acyltransferase (GPAT), 1-Acylglycerol-3-phosphate Acyltransferase (AGPAT), and Diacylglycerol Acyltransferase (DGAT), which are present on ER and mitochondrial membranes. The formation of CEs depends on Acyl CoA Cholesterol Acyltransferase, an ER protein which would again be affected by ER architecture (Henne et al., 2025; Smolic et al., 2021). Thus, a drop in LD density or size may result from one or more of the following. (i) A lack of ceramide production, which affects ER structure and thus influences LD formation. Earlier studies have suggested that ceramide depletion leads to ER expansion and compromised plasma membrane integrity (Rao et al., 2014; Rao et al., 2007; Zhu et al., 2025). (ii) Increase in various ceramide intermediates within the ER, and subsequent diversion of the lipid flux to other metabolic pathways (iii) Limited production of complex sphingolipids due to a reduction of ceramide either being produced or being transferred from the ER (iv) Lipolysis – a drop in lipid density and size could be a result of an upswing in LD utilization for membrane maintenance. One possible reason for the requirement for extra membranes would be Glial roles in phagocytosis and axonal ensheathment, which would be affected in the absence of sphingolipids (Ghosh et al., 2013; Kunduri et al., 2025; Theisen et al., 2025; Vaughen et al., 2022). (v) A modulation in the dynamics of MCS between ER and mitochondria or ER and Golgi. Experiments that modulate VAPB and thus affect MCS dynamics (Alpy et al., 2013; Dong et al., 2016; Kalarikkal et al., 2024; Obara et al., 2024) have been shown to affect cellular physiology. In our study, we leave the VAPB tether intact, perturbing only its interaction with CERT (Mishra et al., 2024), and this specific disruption modulates LD dynamics with fewer LDs, but does not affect size.

Thus, in summary, our data points to a direct link between ceramide production and/or its transfer to a vulnerability in the motor neuronal circuit (Fig. 7) that relates to climbing. Glia are more sensitive to disruptions in ceramide synthesis and transport, with impairments leading to adult motor dysfunction. Glial cells appear to be major players in maintaining adult motor function, with LDs serving as a useful readout of glial ceramide status and markers of motor dysfunction.

## Materials and Methods

### *Drosophila* husbandry, fly stocks and reagents

All the experimental stocks were maintained and crossed on a standard corn agar media in 12 hr Light: 12hr Dark cycle at 25 ℃ and F1 male progeny were used for experiments, which were performed at 25℃. *w1118* is used as wild-type control in the experiments. *dCERT^1^ (CERT^Δ^)* was a kind gift from Jairaj Acharya (Rao et al., 2007). Genomic *CERT^WT^* and *CERT^SL^* lines were acquired from Charlotte Gehin & Giovanni D’Angelo (Gehin et al., 2023). *UASp-CERT^WT^* and *UASp-CERT^EAAD^* transgenic lines were generated in-house. The UAS-Gal4 system was used for OE and KD experiments (Andrea H. Brand, 1993). *repo-Gal4* was a kind gift from Dr. Bradley Jones. *elav-Gal4* has been described in (Ratnaparkhi et al., 2008). For *CERT* KD, Valium 20 Trip RNAi line were used unless specified. Bloomington fly Stocks: *w1118* (BL_3605), *Df(3L)Exel6112* (BL_7591), *tub-Gal4*(BL_5138), *nSyb-Gal4*(BL_51635), *CERT^RNAi^* (BL_60080), *CERT^RNAi^* (BL_35579), *spt-I^RNAi^* (BL_55685), *lace^RNAi^* (BL_51475), *Kdsr^RNAi^* (BL_64494), *schlank^RNAi^* (BL_29340), *ifc^RNAi^* (BL_32514) were used for experiments.

### Cloning and generation of *CERT* transgenic lines

*CERT* cDNA from *Drosophila* Gold collection (https://dgrc.bio.indiana.edu/) was amplified using *CERT* specific primers. Site specific mutagenesis was used to create FF>AA mutation (306-307 AA position) in the *CERT* DNA fragment. The second-chromosome UAS transgenics for *CERT^WT^ and CERT^EAAD^* were generated by targeted insertion at attP40 using the pUASp-attB vector obtained from the *Drosophila* Genomics Resource Centre (DGRC, #1358). Transgenics were balanced, screened using min-white as a selectable marker and validated using qPCR. Following primers (5’-> 3’ direction) were used: *CERT* cDNA-*ATGGACACGAACGCTGGGGAG* (forward), *TCAGAACATGATCGGCTTGTCCTT* (reversed). For cloning in pUASp-attB vector: *GGATCAGATCCGCGGCCGCATGGACACGAACGCTGGGGAG* (forward), *CCACTAGTGGCCTATGCGGCCGCTCAGAACATGATCGGCTTGTC* (reversed) and for pUASp-CERT DNA sequencing: *GGCAAGGGTCGAGTCGATAG* (forward) and *AGGTTTAACCAGGGGATGCT* (reversed), for FF>AA DNA sequencing: *GGACCAAAGCAAAAAGCTCAA* (forward) and *GCGATCGATCTCCGGCCACAG* (reversed).

### Real Time PCR

mRNA was extracted from 10-day-old male flies (10 flies per genotype) using the Abcam manual extraction protocol. A total of 1µg RNA was used for cDNA synthesis using the High-Capacity cDNA reverse transcriptase Kit (4368814) by Invitrogen. Then qPCR was performed using gene specific primers along with iTaq Universal SyBr Green Supermix (Biorad, #1725124) in qTOWER3 778 Series Real Time PCR 779 Thermal Cycler. The relative fold change for each genotype was calculated by normalizing to housekeeping gene *rp49*. Experiment was performed for three replicates per genotype. *CERT^RNAi(V20)^* (0.24), *CERT^WT^* (14.6), *CERT^EAAD^* (25.7) compared to control. Following primer pairs (5’->3’) were used: *rp49: GACGCTTCAAGGGACAGTATC* (forward); *AAACGCGGTTCTGCATGAG* (reverse), *CERT: ATCGATTTCAAGGGCGAATCCATC* (forward); *TCCTTGAGCTTTTTGCTTTGGTCC* (reverse), FFAT: *ACATTGCCCGAGGATGAGTTCTTC*(forward) and *GCGATCGATCTCCGGCCACAG* (reverse).

### Lipid Droplet staining for adult brains

For adult brains, flies of the required genotype were anaesthetized on ice and brains were dissected in ice-cold 1X PBS. Fixation was done for 30 minutes with 4% PFA (0.5% Triton-x-100) followed by two immediate washes of 1XPBS and then 3^rd^ wash of 15 minutes with 1 XPBS at room temperature. For LD staining, 1:1000 BODIPY^TM^ 493/503 (1mg/ml, Invitrogen, D3922) along with 1:1000 Hoescht (10mg/ml, Invitrogen) in 1XPBS was added to the brains and incubated on a rocker for 30 minutes at room temperature under dark conditions. Then, the solution was removed, and the brains were once rinsed with 1X PBS. Bridge mounting was done for adult brains using slowfade mounting reagent (Vectashield S36937, Invitrogen), and imaging was done within 24 hours of sample preparation.

### Microscope imaging and analysis

Mounted samples were imaged on Leica SP8 with 63X objective lens. Z-stacks were acquired with zoom factor of 0.75X unless specified with 1µm interval at 16-bit depth. Other settings were kept constant across experiments with changes in laser power and gain for individual experiments. For image analysis, Fiji software was used where manual protocol was made for analysing lipid droplets. Firstly, mean intensity value of each slice was subtracted from that slice of the stack using a macro code. For adult brains, cells surrounding subesophageal zone (SEZ) region with five ROIs were taken for LD analysis. A macro code is made which automatically process the ROI and analyse lipid droplets using intensity and size as thresholding parameters. Briefly, an arbitrary constant value was subtracted to reduce the noise and then mean 3D filter was used. Automatic thresholding was done using Otsu method for each slice of stack and then processed through 3D object counter for analysing the droplets with size filtration of 8-1000. Macro code was able to store each ROI data into excel file where number and volume of droplets can be used for plotting the data. For lipid droplet density, the total number of droplets in each ROI was divided by the volume of the ROI based on the nuclei stack (number of droplets per µm^3^). Lipid droplet size was plotted in the form of a scattered dot plot by taking the volume of all droplets in the ROIs from all brain samples of each genotype into consideration.

### Climbing Assay

Climbing activity of each genotype was observed using the startle-induced negative geotaxis assay as described (Tendulkar et al., 2022). Three biological replicates of each genotype with 25-30 age-matched male flies were raised at 25 ℃. Each set was transferred to 250ml glass cylinder and allowed to acclimatize for 5 minutes. Then, cylinder was tapped three times to startle the flies and the flies were allowed to climb for 1 minute. After 1 minute, flies were counted based on three classes, non-climbers, poor climbers (0-80ml, 7.5cm) and good climbers (above 80ml or 7.5cm). These classes were given scores of 0, 1 and 2. A 3-minute rest period was given between sets, and the set was repeated twice more. The same protocol was followed for biological replicates and rest of the genotypes. The experiment was done every five days from eclosion, upto 30 days. CO_2_ was used to anaesthetise the flies after 3 trials, and they were transferred to fresh media vials. Climbing Index was used to represent the motor activity of flies, where the given score was multiplied by the number of flies in that range, and the sum of all three values was averaged with the 3 trial values. Then, the number was divided by the total flies taken for that set.

### Statistical tests

Two way ANOVA was used for climbing assays followed by Tukey’s test for multiple comparisons. For LD density, ordinary one-way ANOVA was used and then multiple comparisons were done using Tukey’s test. For LD size, one-way ANOVA was performed using the Kruskal-Wallis test followed by Dunn’s test for multiple comparison correction. Graphpad Prism 10.4.1(532) was used for statistical tests, and BioRender and Microsoft PowerPoint for schematic diagrams.

## Data & Resource availability

Raw imaging data will be shared on acceptance of the manuscript, as per journal policy.

## Contributions

LG and GSR conceptualised the project. LG executed the experiments, with KH contributing to the ceramide screen. LG wrote the initial draft with inputs from GSR. SD and AR generated the *CERT* transgenic lines. GSR was responsible for funding acquisition, supervision and project management.

## Acknowledgements

We thank: Bloomington *Drosophila* Stock Centre (BDSC), supported by NIH grant P40OD018537, for fly stocks; Drosophila Genomics Resource Centre, supported by NIH grant 2P40OD01094, for the Gold cDNA library; Snehal Patil and Yashwant Pawar for fly media and stock maintenance. The fly facility at the National Centre for Biological Sciences (NCBS), Bangalore, for injecting *CERT* constructs into *Drosophila* embryos; Microscopy facility at IISER Pune, managed by Dr Santosh Podder and Vijay Vitthal.

## References

Agrawal, I., Lim, Y. S., Ng, S. Y. and Ling, S. C. (2022). Deciphering lipid dysregulation in ALS: from mechanisms to translational medicine. Transl Neurodegener 11, 48.

Alessenko, A. V. and Albi, E. (2020). Exploring Sphingolipid Implications in Neurodegeneration. Front Neurol 11, 437.

Alpy, F., Rousseau, A., Schwab, Y., Legueux, F., Stoll, I., Wendling, C., Spiegelhalter, C., Kessler, P., Mathelin, C., Rio, M. C. et al. (2013). STARD3 or STARD3NL and VAP form a novel molecular tether between late endosomes and the ER. J Cell Sci 126, 5500–12.

Andrea H. Brand, N. P. (1993). Targeted gene expression as a means of altering cell fates and generating dominant phenotypes. Development 118, 401–415.

Arboleda, G., Morales, L. C., Benitez, B. and Arboleda, H. (2009). Regulation of ceramide-induced neuronal death: cell metabolism meets neurodegeneration. Brain Res Rev 59, 333–46.

Ayub, M., Jin, H. K. and Bae, J. S. (2021). Novelty of Sphingolipids in the Central Nervous System Physiology and Disease: Focusing on the Sphingolipid Hypothesis of Neuroinflammation and Neurodegeneration. Int J Mol Sci 22.

Barber, C. N. and Raben, D. M. (2019). Lipid Metabolism Crosstalk in the Brain: Glia and Neurons. Front Cell Neurosci 13, 212.

Bauer, R., Voelzmann, A., Breiden, B., Schepers, U., Farwanah, H., Hahn, I., Eckardt, F., Sandhoff, K. and Hoch, M. (2009). Schlank, a member of the ceramide synthase family controls growth and body fat in Drosophila. EMBO J 28, 3706–16.

Bikman, B. T. and Summers, S. A. (2011). Ceramides as modulators of cellular and whole-body metabolism. J Clin Invest 121, 4222–30.

Blair, K., Martinez-Serra, R., Gosset, P., Martin-Guerrero, S. M., Morotz, G. M., Atherton, J., Mitchell, J. C., Markovinovic, A. and Miller, C. C. J. (2025). Structural and functional studies of the VAPB-PTPIP51 ER-mitochondria tethering proteins in neurodegenerative diseases. Acta Neuropathol Commun 13, 49.

Bohnert, M. and Schrul, B. (2024). Lipid droplets in health and disease. FEBS Lett 598, 1113–1115.

Borgese, N., Iacomino, N., Colombo, S. F. and Navone, F. (2021a). The Link between VAPB Loss of Function and Amyotrophic Lateral Sclerosis. Cells 10.

Borgese, N., Navone, F., Nukina, N. and Yamanaka, T. (2021b). Mutant VAPB: Culprit or Innocent Bystander of Amyotrophic Lateral Sclerosis? Contact 4.

Byrns, C. N., Perlegos, A. E., Miller, K. N., Jin, Z., Carranza, F. R., Manchandra, P., Beveridge, C. H., Randolph, C. E., Chaluvadi, V. S., Zhang, S. L., et al. (2024). Senescent glia link mitochondrial dysfunction and lipid accumulation. Nature 630, 475–483.

Chaplot, K., Pimpale, L., Ramalingam, B., Deivasigamani, S., Kamat, S. S. and Ratnaparkhi, G. S. (2019). SOD1 activity threshold and TOR signalling modulate VAP(P58S) aggregation via reactive oxygen species-induced proteasomal degradation in a Drosophila model of amyotrophic lateral sclerosis. Dis Model Mech 12.

Chen, X., Li, J., Gao, Z., Yang, Y., Kuang, W., Dong, Y., Chua, G. H., Huang, X., Jiang, B., Tian, H. et al. (2022). Endogenous ceramide phosphoethanolamine modulates circadian rhythm via neural-glial coupling in Drosophila. Natl Sci Rev 9, nwac148.

Cockcroft, S. and Raghu, P. (2018). Phospholipid transport protein function at organelle contact sites. Curr Opin Cell Biol 53, 52–60.

Crivelli, S. M., Giovagnoni, C., Zhu, Z., Tripathi, P., Elsherbini, A., Quadri, Z., Pu, J., Zhang, L., Ferko, B., Berkes, D. et al. (2022). Function of ceramide transfer protein for biogenesis and sphingolipid composition of extracellular vesicles. J Extracell Vesicles 11, e12233.

Cutler, R. G., Pedersen, W. A., Camandola, S., Rothstein, J. D. and Mattson, M. P. (2002). Evidence that accumulation of ceramides and cholesterol esters mediates oxidative stress-induced death of motor neurons in amyotrophic lateral sclerosis. Ann Neurol 52, 448–57.

Deivasigamani, S., Verma, H. K., Ueda, R., Ratnaparkhi, A. and Ratnaparkhi, G. S. (2014). A genetic screen identifies Tor as an interactor of VAPB in a Drosophila model of amyotrophic lateral sclerosis. Biol Open 3, 1127–38.

Dong, R., Saheki, Y., Swarup, S., Lucast, L., Harper, J. W. and De Camilli, P. (2016). Endosome-ER Contacts Control Actin Nucleation and Retromer Function through VAP-Dependent Regulation of PI4P. Cell 166, 408–423.

Fullam, T. and Statland, J. (2021). Upper Motor Neuron Disorders: Primary Lateral Sclerosis, Upper Motor Neuron Dominant Amyotrophic Lateral Sclerosis, and Hereditary Spastic Paraplegia. Brain Sci 11.

Gehin, C., Lone, M. A., Lee, W., Capolupo, L., Ho, S., Adeyemi, A. M., Gerkes, E. H., Stegmann, A. P., Lopez-Martin, E., Bermejo-Sanchez, E. et al. (2023). CERT1 mutations perturb human development by disrupting sphingolipid homeostasis. J Clin Invest 133.

Ghosh, A., Kling, T., Snaidero, N., Sampaio, J. L., Shevchenko, A., Gras, H., Geurten, B., Gopfert, M. C., Schulz, J. B., Voigt, A. et al. (2013). A global in vivo Drosophila RNAi screen identifies a key role of ceramide phosphoethanolamine for glial ensheathment of axons. PLoS Genet 9, e1003980.

Goodman, L. D., Ralhan, I., Li, X., Lu, S., Moulton, M. J., Park, Y. J., Zhao, P., Kanca, O., Ghaderpour Taleghani, Z. S., Jacquemyn, J. et al. (2024). Tau is required for glial lipid droplet formation and resistance to neuronal oxidative stress. Nat Neurosci 27, 1918–1933.

Goyal, G., Zheng, J., Adam, E., Steffes, G., Jain, M., Klavins, K. and Hummel, T. (2019). Sphingolipid-dependent Dscam sorting regulates axon segregation. Nat Commun 10, 813.

Gunay, A., Shin, H. H., Gozutok, O., Gautam, M. and Ozdinler, P. H. (2021). Importance of lipids for upper motor neuron health and disease. Semin Cell Dev Biol 112, 92–104.

Hanada, K. (2014). Co-evolution of sphingomyelin and the ceramide transport protein CERT. Biochim Biophys Acta 1841, 704–19.

Hebbar, S., Sahoo, I., Matysik, A., Argudo Garcia, I., Osborne, K. A., Papan, C., Torta, F., Narayanaswamy, P., Fun, X. H., Wenk, M. R. et al. (2015). Ceramides And Stress Signalling Intersect With Autophagic Defects In Neurodegenerative Drosophila blue cheese (bchs) Mutants. Sci Rep 5, 15926.

Henne, W. M., Reynolds, E. and Prinz, W. A. (2025). Lipid droplets: Open questions and conceptual advances around a unique organelle. J Cell Biol 224.

Henriques, A., Croixmarie, V., Bouscary, A., Mosbach, A., Keime, C., Boursier-Neyret, C., Walter, B., Spedding, M. and Loeffler, J. P. (2017). Sphingolipid Metabolism Is Dysregulated at Transcriptomic and Metabolic Levels in the Spinal Cord of an Animal Model of Amyotrophic Lateral Sclerosis. Front Mol Neurosci 10, 433.

Hussain, G., Wang, J., Rasul, A., Anwar, H., Imran, A., Qasim, M., Zafar, S., Kamran, S. K. S., Razzaq, A., Aziz, N. et al. (2019). Role of cholesterol and sphingolipids in brain development and neurological diseases. Lipids Health Dis 18, 26.

Ilieva, H., Polymenidou, M. and Cleveland, D. W. (2009). Non-cell autonomous toxicity in neurodegenerative disorders: ALS and beyond. J Cell Biol 187, 761–72.

Islam, M. N., Mandal, N., Khanom, N. I., Spivey, A. L., Patel, A. S., Ahmed, T., Islam, R. and Bhuiyan, M. I. H. (2025). Sphingolipid Signaling and Metabolism in Neuronal and Glial Cells: Implications for Cerebrovascular and Neurodegenerative Disorders. Aging Dis.

James, C. and Kehlenbach, R. H. (2021). The Interactome of the VAP Family of Proteins: An Overview. Cells 10.

Jeon, S., Scorletti, E., Dempsey, J., Buyco, D., Lin, C., Saiman, Y., Bayen, S., Harkin, J., Martin, J., Hooks, R. et al. (2023). Ceramide synthase 6 (CerS6) is upregulated in alcohol-associated liver disease and exhibits sex-based differences in the regulation of energy homeostasis and lipid droplet accumulation. Mol Metab 78, 101804.

Jung, W. H., Liu, C. C., Yu, Y. L., Chang, Y. C., Lien, W. Y., Chao, H. C., Huang, S. Y., Kuo, C. H., Ho, H. C. and Chan, C. C. (2017). Lipophagy prevents activity-dependent neurodegeneration due to dihydroceramide accumulation in vivo. EMBO Rep 18, 1150–1165.

Kalarikkal, M., Saikia, R., Oliveira, L., Bhorkar, Y., Lonare, A., Varshney, P., Dhamale, P., Majumdar, A. and Joseph, J. (2024). Nup358 restricts ER-mitochondria connectivity by modulating mTORC2/Akt/GSK3beta signalling. EMBO Rep 25, 4226–4251.

Kamemura, K., Chen, C. A., Okumura, M., Miura, M. and Chihara, T. (2021). Amyotrophic lateral sclerosis-associated Vap33 is required for maintaining neuronal dendrite morphology and organelle distribution in Drosophila. Genes Cells 26, 230–239.

Kawano, M., Kumagai, K., Nishijima, M. and Hanada, K. (2006). Efficient trafficking of ceramide from the endoplasmic reticulum to the Golgi apparatus requires a VAMP-associated protein-interacting FFAT motif of CERT. J Biol Chem 281, 30279–88.

Kentaro Hanada, K. K., Satoshi Yasuda, Yukiko Miura, Miyuki Kawano, Masayoshi Fukasawa & Masahiro Nishijima. (2003). Molecular machinery for non-vesicular trafficking of ceramide. Nature 426, 803–809.

Kolter, T. and Sandhoff, K. (1999). Sphingolipids-Their Metabolic Pathways and the Pathobiochemistry of Neurodegenerative Diseases. Angew Chem Int Ed Engl 38, 1532–1568.

Kors, S., Costello, J. L. and Schrader, M. (2022). VAP Proteins - From Organelle Tethers to Pathogenic Host Interactors and Their Role in Neuronal Disease. Front Cell Dev Biol 10, 895856.

Kovacs, D., Gay, A. S., Debayle, D., Abelanet, S., Patel, A., Mesmin, B., Luton, F. and Antonny, B. (2024). Lipid exchange at ER-trans-Golgi contact sites governs polarized cargo sorting. J Cell Biol 223.

Kraut, R. (2011). Roles of sphingolipids in Drosophila development and disease. J Neurochem 116, 764–78.

Kumagai, K. and Hanada, K. (2019). Structure, functions and regulation of CERT, a lipid-transfer protein for the delivery of ceramide at the ER-Golgi membrane contact sites. FEBS Lett 593, 2366–2377.

Kunduri, G., Godenschwege, T. A., Sankey, K., Abhilasha, K. V., Acharya, U. R. and Acharya, J. K. (2025). Light and temperature sensitive seizures are regulated by spatially distinct cortex glial populations in the central nervous system. bioRxiv.

Lee, J. A., Hall, B., Allsop, J., Alqarni, R. and Allen, S. P. (2021). Lipid metabolism in astrocytic structure and function. Semin Cell Dev Biol 112, 123–136.

Liping Wang, G. L., Zhongyuan Zuo, Yarong Li, Seul Kee Byeon, Akhilesh Pandey, Hugo J. Bellen. (2022). Neuronal activity induces glucosylceramide that is secreted via exosomes for lysosomal degradation in glia. Science Advances.

Liu, J., Luo, X., Chen, X. and Shang, H. (2020). Lipid Profile in Patients With Amyotrophic Lateral Sclerosis: A Systematic Review and Meta-Analysis. Front Neurol 11, 567753.

Liu, L., MacKenzie, K. R., Putluri, N., Maletic-Savatic, M. and Bellen, H. J. (2017). The Glia-Neuron Lactate Shuttle and Elevated ROS Promote Lipid Synthesis in Neurons and Lipid Droplet Accumulation in Glia via APOE/D. Cell Metab 26, 719–737 e6.

Liu, L., Zhang, K., Sandoval, H., Yamamoto, S., Jaiswal, M., Sanz, E., Li, Z., Hui, J., Graham, B. H., Quintana, A. et al. (2015). Glial lipid droplets and ROS induced by mitochondrial defects promote neurodegeneration. Cell 160, 177–90.

McCluskey, G., Donaghy, C., Morrison, K. E., McConville, J., Duddy, W. and Duguez, S. (2022). The Role of Sphingomyelin and Ceramide in Motor Neuron Diseases. J Pers Med 12.

McInnis, J. J., Sood, D., Guo, L., Dufault, M. R., Garcia, M., Passaro, R., Gao, G., Zhang, B. and Dodge, J. C. (2024). Unravelling neuronal and glial differences in ceramide composition, synthesis, and sensitivity to toxicity. Commun Biol 7, 1597.

Mencarelli, C. and Martinez-Martinez, P. (2013). Ceramide function in the brain: when a slight tilt is enough. Cell Mol Life Sci 70, 181–203.

Mesmin, B., Kovacs, D. and D’Angelo, G. (2019). Lipid exchange and signaling at ER-Golgi contact sites. Curr Opin Cell Biol 57, 8–15.

Mishra, S., Manohar, V., Chandel, S., Manoj, T., Bhattacharya, S., Hegde, N., Nath, V. R., Krishnan, H., Wendling, C., Di Mattia, T. et al. (2024). A genetic screen to uncover mechanisms underlying lipid transfer protein function at membrane contact sites. Life Sci Alliance 7.

Moustaqim-Barrette, A., Lin, Y. Q., Pradhan, S., Neely, G. G., Bellen, H. J. and Tsuda, H. (2014). The amyotrophic lateral sclerosis 8 protein, VAP, is required for ER protein quality control. Hum Mol Genet 23, 1975–89.

Murphy, S. E. and Levine, T. P. (2016). VAP, a Versatile Access Point for the Endoplasmic Reticulum: Review and analysis of FFAT-like motifs in the VAPome. Biochim Biophys Acta 1861, 952–961.

Neefjes, J. and Cabukusta, B. (2021). What the VAP: The Expanded VAP Family of Proteins Interacting With FFAT and FFAT-Related Motifs for Interorganellar Contact. Contact *(Thousand Oaks)* 4, 25152564211012246.

Nishimura, A. L., Mitne-Neto, M., Silva, H. C., Richieri-Costa, A., Middleton, S., Cascio, D., Kok, F., Oliveira, J. R., Gillingwater, T., Webb, J. et al. (2004). A mutation in the vesicle-trafficking protein VAPB causes late-onset spinal muscular atrophy and amyotrophic lateral sclerosis. Am J Hum Genet 75, 822–31.

Nowakowski, T. J., Nano, P. R., Matho, K. S., Chen, X., Corrigan, E. K., Ding, W., Gao, Y., Heffel, M., Jayakumar, J., Kaplan, H. S. et al. (2025). The new frontier in understanding human and mammalian brain development. Nature 647, 51–59.

Obara, C. J., Nixon-Abell, J., Moore, A. S., Riccio, F., Hoffman, D. P., Shtengel, G., Xu, C. S., Schaefer, K., Pasolli, H. A., Masson, J. B. et al. (2024). Motion of VAPB molecules reveals ER-mitochondria contact site subdomains. Nature 626, 169–176.

Oswald, M. C., West, R. J., Lloyd-Evans, E. and Sweeney, S. T. (2015). Identification of dietary alanine toxicity and trafficking dysfunction in a Drosophila model of hereditary sensory and autonomic neuropathy type 1. Hum Mol Genet 24, 6899–909.

Pan, X., Dutta, D., Lu, S. and Bellen, H. J. (2023). Sphingolipids in neurodegenerative diseases. Front Neurosci 17, 1137893.

Pan, Y., Li, J., Lin, P., Wan, L., Qu, Y., Cao, L. and Wang, L. (2024). A review of the mechanisms of abnormal ceramide metabolism in type 2 diabetes mellitus, Alzheimer’s disease, and their co-morbidities. Front Pharmacol 15, 1348410.

Pant, D. C., Aguilera-Albesa, S. and Pujol, A. (2020). Ceramide signalling in inherited and multifactorial brain metabolic diseases. Neurobiol Dis 143, 105014.

Pant, D. C., Dorboz, I., Schluter, A., Fourcade, S., Launay, N., Joya, J., Aguilera-Albesa, S., Yoldi, M. E., Casasnovas, C., Willis, M. J. et al. (2019). Loss of the sphingolipid desaturase DEGS1 causes hypomyelinating leukodystrophy. J Clin Invest 129, 1240–1256.

Pathak, D., Mehendale, N., Singh, S., Mallik, R. and Kamat, S. S. (2018). Lipidomics Suggests a New Role for Ceramide Synthase in Phagocytosis. ACS Chem Biol 13, 2280–2287.

Peng, Y. F., Chen, S. J., Li, J. L., Lin, C. H. and Kuo, C. H. (2025). Dysregulated metabolism of ceramides and glycosphingolipids in Parkinson’s disease. J Lipid Res 67, 100955.

Pennetta, G. and Welte, M. A. (2018). Emerging Links between Lipid Droplets and Motor Neuron Diseases. Dev Cell 45, 427–432.

Peretti, D., Dahan, N., Shimoni, E., Hirschberg, K. and Lev, S. (2008). Coordinated lipid transfer between the endoplasmic reticulum and the Golgi complex requires the VAP proteins and is essential for Golgi-mediated transport. Mol Biol Cell 19, 3871–84.

Perry, R. J. and Ridgway, N. D. (2006). Oxysterol-binding protein and vesicle-associated membrane protein-associated protein are required for sterol-dependent activation of the ceramide transport protein. Mol Biol Cell 17, 2604–16.

Rachakonda, V., Pan Th Fau - Le, W. D. and Le, W. D. (2004). Biomarkers of neurodegenerative disorders: how good are they? Cell research 14, 349–360.

Rao, R. P., Scheffer, L., Srideshikan, S. M., Parthibane, V., Kosakowska-Cholody, T., Masood, M. A., Nagashima, K., Gudla, P., Lockett, S., Acharya, U. et al. (2014). Ceramide transfer protein deficiency compromises organelle function and leads to senescence in primary cells. PLoS One 9, e92142.

Rao, R. P., Yuan, C., Allegood, J. C., Rawat, S. S., Edwards, M. B., Wang, X., Merrill, A. H., Jr., Acharya, U. and Acharya, J. K. (2007). Ceramide transfer protein function is essential for normal oxidative stress response and lifespan. Proc Natl Acad Sci U S A 104, 11364–9.

Ratnaparkhi, A., Lawless, G. M., Schweizer, F. E., Golshani, P. and Jackson, G. R. (2008). A Drosophila model of ALS: human ALS-associated mutation in VAP33A suggests a dominant negative mechanism. PLoS One 3, e2334.

Robles-Martinez, L., Morin, K. H. and Nikolova-Karakashian, M. (2025). Ceramide homeostasis in hepatic lipid droplets. Biochemical Society Transactions 53, 509–518.

Schmitt, F., Hussain, G., Dupuis, L., Loeffler, J. P. and Henriques, A. (2014). A plural role for lipids in motor neuron diseases: energy, signaling and structure. Front Cell Neurosci 8, 25.

Senkal, C. E., Salama, M. F., Snider, A. J., Allopenna, J. J., Rana, N. A., Koller, A., Hannun, Y. A. and Obeid, L. M. (2017). Ceramide Is Metabolized to Acylceramide and Stored in Lipid Droplets. Cell Metab 25, 686–697.

Slee, J. A. and Levine, T. P. (2019). Systematic prediction of FFAT motifs across eukaryote proteomes identifies nucleolar and eisosome proteins with the predicted capacity to form bridges to the endoplasmic reticulum. Contact (*Thousand Oaks*) 2, 1–21.

Smolic, T., Zorec, R. and Vardjan, N. (2021). Pathophysiology of Lipid Droplets in Neuroglia. Antioxidants (*Basel*) 11.

Sociale, M., Wulf, A. L., Breiden, B., Klee, K., Thielisch, M., Eckardt, F., Sellin, J., Bulow, M. H., Lobbert, S., Weinstock, N. et al. (2018). Ceramide Synthase Schlank Is a Transcriptional Regulator Adapting Gene Expression to Energy Requirements. Cell Rep 22, 967–978.

Tang, Y., Majewska, M., Less, B., Mehmeti, I., Wollnitzke, P., Semleit, N., Levkau, B., Saba, J. D., van Echten-Deckert, G. and Gurgul-Convey, E. (2024). The fate of intracellular S1P regulates lipid droplet turnover and lipotoxicity in pancreatic beta-cells. J Lipid Res 65, 100587.

Tendulkar, S., Hegde, S., Garg, L., Thulasidharan, A., Kaduskar, B., Ratnaparkhi, A. and Ratnaparkhi, G. S. (2022). Caspar, an adapter for VAPB and TER94, modulates the progression of ALS8 by regulating IMD/NFkappaB-mediated glial inflammation in a Drosophila model of human disease. Hum Mol Genet 31, 2857–2875.

Theisen, E. K., Rivas-Serna, I. M., Lee, R. J., Jay, T. R., Kunduri, G., Nguyen, T. T., Mazurak, V., Clandinin, M. T., Clandinin, T. R. and Vaughen, J. P. (2025). Glia phagocytose neuronal sphingolipids to infiltrate developing synapses. bioRxiv.

Thulasidharan, A., Garg, L., Tendulkar, S. and Ratnaparkhi, G. S. (2024). Age-dependent dynamics of neuronal VAPB(ALS) inclusions in the adult brain. Neurobiol Dis 196, 106517.

Toprak, U., Hegedus, D., Dogan, C. and Guney, G. (2020). A journey into the world of insect lipid metabolism. Arch Insect Biochem Physiol 104, e21682.

Tracey, T. J., Kirk, S. E., Steyn, F. J. and Ngo, S. T. (2021). The role of lipids in the central nervous system and their pathological implications in amyotrophic lateral sclerosis. Semin Cell Dev Biol 112, 69–81.

Uranbileg, B., Sakai, E., Kubota, M., Isago, H., Sumitani, M., Yatomi, Y. and Kurano, M. (2024). Development of an advanced liquid chromatography-tandem mass spectrometry measurement system for simultaneous sphingolipid analysis. Sci Rep 14, 5699.

Vacaru, A. M., Tafesse, F. G., Ternes, P., Kondylis, V., Hermansson, M., Brouwers, J. F., Somerharju, P., Rabouille, C. and Holthuis, J. C. (2009). Sphingomyelin synthase-related protein SMSr controls ceramide homeostasis in the ER. J Cell Biol 185, 1013–27.

Vaughen, J. P., Theisen, E., Rivas-Serna, I. M., Berger, A. B., Kalakuntla, P., Anreiter, I., Mazurak, V. C., Rodriguez, T. P., Mast, J. D., Hartl, T. et al. (2022). Glial control of sphingolipid levels sculpts diurnal remodeling in a circadian circuit. Neuron 110, 3186–3205 e7.

Verma, R., Sharma, P., Sharma, V. and Singh, T. G. (2025). Modulating lipid droplet dynamics in neurodegeneration: an emerging area of molecular pharmacology. Mol Biol Rep 52, 277.

Voelzmann, A. and Bauer, R. (2010). Ceramide synthases in mammalians, worms, and insects: emerging schemes. Biomol Concepts 1, 411–22.

Vos, M., Dulovic-Mahlow, M., Mandik, F., Frese, L., Kanana, Y., Haissatou Diaw, S., Depperschmidt, J., Bohm, C., Rohr, J., Lohnau, T. et al. (2021). Ceramide accumulation induces mitophagy and impairs beta-oxidation in PINK1 deficiency. Proc Natl Acad Sci U S A 118.

Walther, T. C. and Farese, R. V., Jr. (2012). Lipid droplets and cellular lipid metabolism. Annu Rev Biochem 81, 687–714.

Wang, X., Rao, R. P., Kosakowska-Cholody, T., Masood, M. A., Southon, E., Zhang, H., Berthet, C., Nagashim, K., Veenstra, T. K., Tessarollo, L. et al. (2009). Mitochondrial degeneration and not apoptosis is the primary cause of embryonic lethality in ceramide transfer protein mutant mice. J Cell Biol 184, 143–58.

Wang, Y., Wang, B., Hou, J., Huo, X., Liu, C., Guan, R., Chen, H., Zhou, Y., Zhang, J., Zhuang, C. et al. (2026). Lipid droplets in astrocytes: Key organelles for CNS homeostasis and disease (Review). Int J Mol Med 57.

Wanikawa, M., Nakamura, H., Emori, S., Hashimoto, N. and Murayama, T. (2020). Accumulation of sphingomyelin in Niemann-Pick disease type C cells disrupts Rab9-dependent vesicular trafficking of cholesterol. J Cell Physiol 235, 2300–2309.

Weber-Boyvat, M., Kentala, H., Peranen, J. and Olkkonen, V. M. (2015). Ligand-dependent localization and function of ORP-VAP complexes at membrane contact sites. Cell Mol Life Sci 72, 1967–87.

Yamaji, T., Kumagai, K., Tomishige, N. and Hanada, K. (2008). Two sphingolipid transfer proteins, CERT and FAPP2: their roles in sphingolipid metabolism. IUBMB Life 60, 511–8.

Yang, D., Wang, X., Zhang, L., Fang, Y., Zheng, Q., Liu, X., Yu, W., Chen, S., Ying, J. and Hua, F. (2022). Lipid metabolism and storage in neuroglia: role in brain development and neurodegenerative diseases. Cell Biosci 12, 106.

Zhang, L., Zhou, Y., Yang, Z., Jiang, L., Yan, X., Zhu, W., Shen, Y., Wang, B., Li, J. and Song, J. (2025). Lipid droplets in central nervous system and functional profiles of brain cells containing lipid droplets in various diseases. J Neuroinflammation 22, 7.

Zhu, Y., Cho, K., Lacin, H., Zhu, Y., DiPaola, J. T., Wilson, B. A., Patti, G. and Skeath, J. B. (2025). Loss of dihydroceramide desaturase drives neurodegeneration by disrupting endoplasmic reticulum and lipid droplet homeostasis in glial cells. Elife 13.

