## Supplementary material for "Glial ceramide orchestrates Lipid Droplet homeostasis and age-dependent motor function in *Drosophila*": Suppl. Files

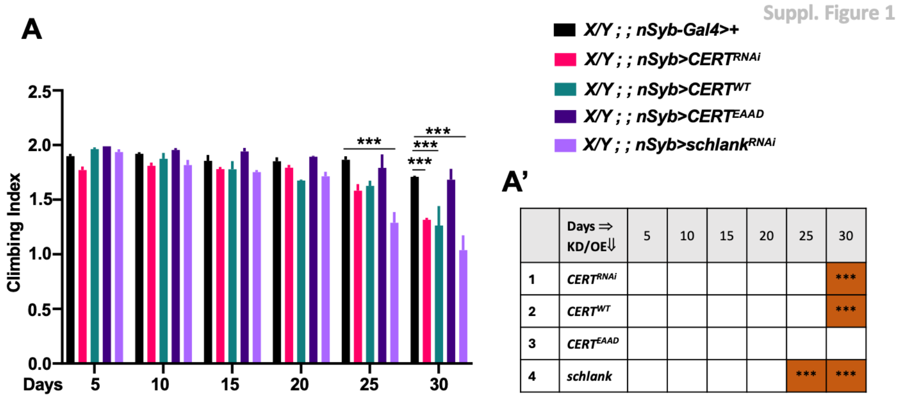


| | Days ⇒ KD/OE⇓ | 5 | 10 | 15 | 20 | 25 | 30 |
|---|---|---|---|---|---|---|---|
| 1 | *CERT*$^{RNAi}$ | | | | | | *** |
| 2 | *CERT*$^{WT}$ | | | | | | *** |
| 3 | *CERT*$^{EAAD}$ | | | | | | |
| 4 | *schlank* | | | | | *** | *** |

**Suppl. Figure 1: Neuronal role of *CERT* in motor function of *Drosophila*.** *CERT* KD and OE using *nSyb-Gal4* showed motor defects after day 25, however, *CERT*$^{EAAD}$ in neurons did not show defects. Also, pan-neuronal *schlank* knockdown led to motor defects at day 25 & 30. (A') Summary table of the gene modulation with statistical significance shown for each day compared to the control. Mean±SEM values are plotted in the graph with p<0.001(***), Climbing Index is represented on Y-axis with number of days after eclosion on X-axis. n=25-30 males, N=3 biological replicates for each genotype.



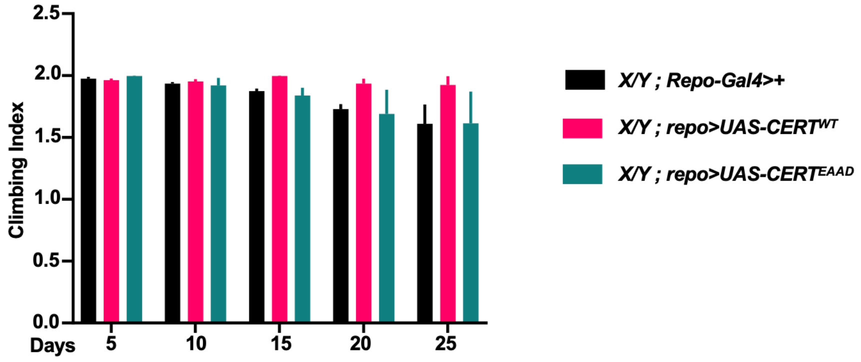


**Suppl. Figure 2: Glial role for *CERT* in motor function of *Drosophila*.** OE of *CERT*$^{EAAD}$ in glia showed no change in motor activity of male flies. Mean±SEM values are plotted in the graph with p<0.001(***), Climbing Index is represented on Y-axis with number of days after eclosion on X-axis. n=25-30 males, N=3 biological replicates for each genotype.

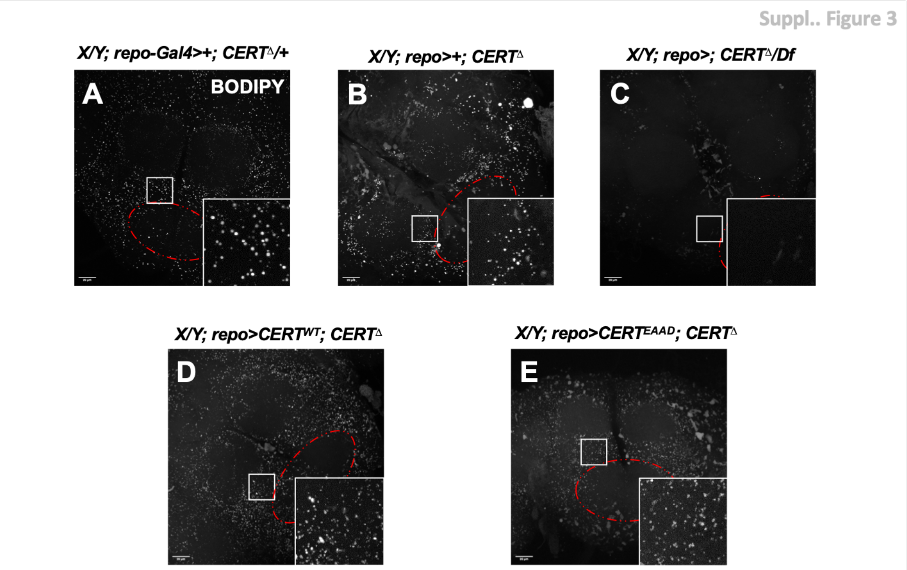


**Suppl. Figure 3: Rescue of LD defects with glial *CERT OE.*** (A-E) represents MIP projections (40% brightness) of BODIPY-stained LDs in the brain of 5-day-old males. *CERT*$^{WT}$ OE (D) in glia of *CERT*$^{\Delta}$ showed LD rescue when compared to (B) and (C). LDs were smaller in (D) than in (A). However, *CERT*$^{EAAD}$ OE rescue leads to fewer and smaller LDs compared to (A) but more LDs compared to (C).
